# Functional Transcriptome Analysis of Bladder Cancer Cell Lines Persistently Infected with Oncolytic Newcastle Disease Virus

**DOI:** 10.1101/2020.12.14.422610

**Authors:** Umar Ahmad, Arcana Thirumorthy, De Ming Chau, Suet Lin Chia, Khatijah Yusoff, Syahril Abdullah, Soon Choy Chan, Abhi Veerakumarasivam

## Abstract

**Background:** Newcastle disease virus (NDV) has been an attractive virotherapy agent that targets various type of human cancers while leaving normal cells unharmed. Wild-type NDV strain AF2240 has been found to persistently infect subpopulation of cancer cells *in vitro*, making the cells less susceptible to NDV-mediated oncolysis. It is proposed that transcriptome profiling of NDV persistently infected bladder cancer cell lines will provide insights to understand such occurrence by identifying specific pathways associated with NDV persistent infection due to transcriptomic dysregulation.

**Results:** Transcriptome profiling revealed a total of 63 and 134 differentially expressed genes (DEGs) from NDV persistently infected TCCSUPPi and EJ28Pi bladder cancer cells relative to their uninfected controls, respectively. Of the 63 DEGs identified for TCCSUPPi cells, 25 DEGs were upregulated (log_2_ fold-change ≥ 0) and 38 DEGs were downregulated (log_2_ fold-change ≤ 0). These genes were significantly enriched in the molecular function of calcium binding (GO:0005509) and DNA-binding transcription repressor activity, RNA polymerase II-specific (GO:0001227) and the enriched important upregulated pathways were mainly heme metabolism, TGF-beta signaling and spermatogenesis. As for EJ28Pi, 55 DEGs were upregulated (log_2_ fold-change ≥ 0) and 79 DEGs were downregulated (log_2_ fold-change ≤ 0). These DEGs resulted in significantly enriched molecular function such as protein domain specific binding (GO:0019904) and RNA polymerase II regulatory region sequence-specific DNA binding (GO:0000977). The enriched important upregulated pathways were allograft rejection, KRAS signaling up and interferon gamma response. Other important pathways that were downregulated in both the NDV-persistently infected cell lines were angiogenesis, apoptosis, and xenobiotic metabolism.

**Conclusion:** The transcriptome profiles (RNA-Seq) of these cell lines suggest that evasion of apoptosis and increase in TGF-beta signaling and interferon gamma response activities are crucial for establishment of NDV persistent infection in bladder cancer cells. Findings from this study provide the molecular basis that warrant further study on how bladder cancer cells acquired NDV persistent infection. Resolving the mechanism of persistent infection will facilitate the application of NDV for more effective treatment of bladder cancer.

## Introduction

Newcastle Disease Virus (NDV) is an oncolytic virus belonging to a promising novel class of virotherapy agent against human cancers while leaving normal cells unharmed[1, 2]. NDV is a member of the new genus *Avulavirus* within the family *Paramyxoviridae* with a non-segmented negative-stranded RNA genome of about 15.9 kb long[3, 4] that contains six genes[5], which encodes for at least eight proteins [6, 7]. The six essential structural proteins: nucleoprotein (NP), phosphoprotein (P), and the large polymerase protein (L) form the nucleocapsid[8, 9] while haemagglutinin-neuraminidase (HN) and fusion protein (F) attached in the lipid bilayer of membrane in the external envelope[10, 11], and inner layer of the virion envelope is formed by the matrix protein (M). The HN protein mediates the binding of virus to cell surface receptors through its haemagglutinating activity and it also displays neuraminidase activity[10, 11]. The F protein is responsible for fusion between the viral envelope and the target membrane. NDV can also be grown in cell culture which causes cell fusion, resulting from the co-expression of the F and HN proteins that are responsible for viral infectivity and virulence[12, 13]. The two additional non-structural proteins, V and W are formed by the RNA editing process during P gene transcription[14]. Studies have also discovered that the V protein plays an important role in preventing interferon responses and apoptosis in chicken cells, but not in human cells. These species-specific effects of V protein make it a determinant of host range restriction[6, 14]. The W protein also play a role in the viral replication and pathogenesis[14].

NDV has been shown to have higher oncolytic specificity against most cancer cell types as compared to normal cells [15]. Once it is inside the cancer cell, the virus replicates and has the propensity to infect the surrounding cancer cells. The virus also stimulates the host anti-cancer immunological system[16, 17]. Different strains of NDV such as 73-T, MTH68, Italian, Ulster, and HUJ have shown to exhibit an oncolytic activity[17–20]. In addition, the oncolytic effects of two Malaysian strains of NDV, AF2240 and V4-UPM, have also been studied *in vitro* and *in vivo* on several human tumour such as breast cancer[2], astrocytoma cell lines[21], and leukemia[22]. However, the wild-type NDV AF2240 has been found to persistently infect subpopulation of colorectal cancer cells that exhibit a smaller plaque morphology and resist NDV-mediated oncolysis[23]. Thus, persistency of infection poses a potential problem in maximizing the efficacy of this virus for the use of cancer treatment. Although, wild-type NDV has been utilized in clinical trials[24–28], the risk of persistent infection still remains unchanged[29]. There is currently no report on the dysregulated transcripts signature of bladder cancer cell lines persistently infected with oncolytic Newcastle disease virus to understand the molecular basis underlying the development of persistent infection. Thus, understanding the mechanisms by which cancer cells acquire persistent infection and reversing these mechanisms will be crucial in ensuring the clinical efficacy of NDV for cancer treatment.

To explore the molecular mechanism that supports the establishment of NDV persistent infection in bladder cancer cell lines, we employed whole transcriptome sequencing (RNA-Seq) on the persistently infected cell lines by next-generation sequencing (NGS) technology to compare gene expression patterns in these cells and that of the untreated control cells to enable identification of dysregulated genes and signaling pathways associated with persistent infection. The findings from this paper suggest that evasion of apoptosis and increase in TGF-beta signaling and interferon gamma response activities are crucial for establishment of NDV persistent infection in bladder cancer cells.

## Results

### Exploratory analysis of the RNA-Seq data

Sample distance was explored to evaluate similarities and dissimilarities between samples based on their log transformed values of transcripts expression, as expected, the RNA-Seq samples are dissimilar from each other as demonstrated by the distance matrix heatmap and hierarchical clustering (Figure 1). To further gain an insight on how different the RNA-Seq samples are, principal component analysis (PCA) was employed [30] and samples were differentiated based on eigenvalues of gene expression (Supplementary Figure 3). As observed, the replicates in each condition showed similarity in the variances with TCCSUPPi versus TCCSUP (control) having less variance (58.6%) and TCCSUPPi_R3 and TCCSUPPi_R2 samples being relatively closer (Supplementary Figure 3A), meaning the genes in both samples may have similar expression patterns. Compared to TCCSUPPi, EJ28Pi versus EJ28 (control) demonstrated to have greater variance (75.1%) (Supplementary Figure 3B) with none of the samples being closer as found in TCCSUPPi samples, suggesting that each sample has a different pattern of gene expression.

**Figure 1:**
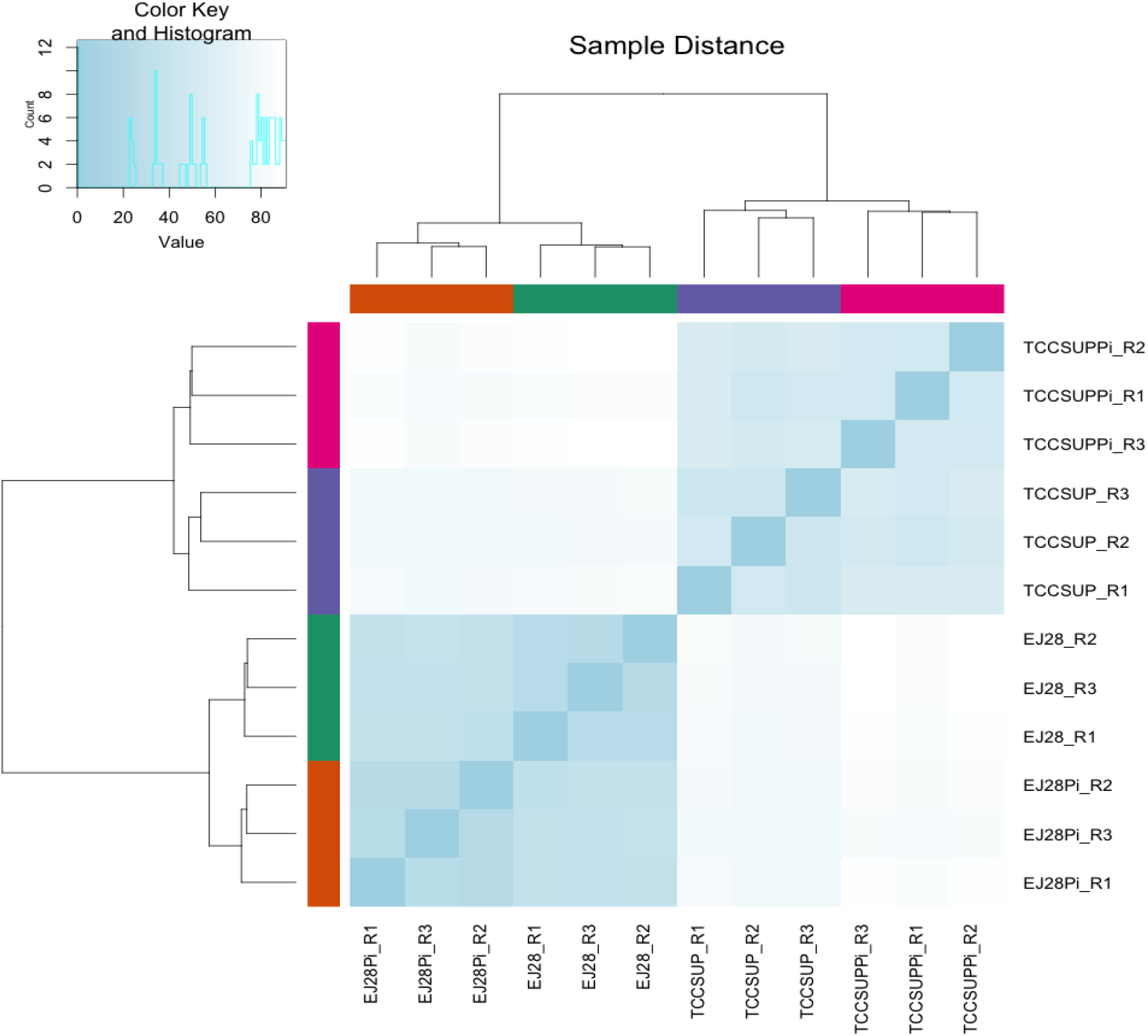
Heatmap clustering of samples based on distance matrix. A heatmap of the distance matrix obtained with DESeq2 package on regularized-logarithm transformed counts to show an overview of similarities and dissimilarities between the RNA-Seq samples. The heatmap showed 2 distinct group of cells (TCCSUP + TCCSUPPi and EJ28Pi + EJ28) and within each group, there is 2 sub-groups of cells (persistently infected cancer cells and control cells).

### Global changes of transcript expression in persistently infected cell lines

To identify the differentially expressed genes (DEGs) that accumulate due to NDV persistent infection in TCCSUP and EJ28 cell lines, gene expression levels were quantified based on fragments per kilobase of exon per million fragments mapped (FPKM) [31] with threshold (q-value) of false discovery rate (FDR) set at 0.05 to judge the statistical significance of the differentially expressed transcripts in the samples. Only transcripts with log2 fold-change ≤ 0 or log2 fold-change ≥ 0 and q-value ≤0.05 are considered differentially expressed genes (DEGs) between persistently infected cells and non-infected control cells. Any genes detected in either non-infected control cells or persistently infected cells based on the above set criteria were considered as uniquely expressed.

Based on FPKM method calculated from DESeq2 package [32], a total of 197 genes were identified in response to NDV persistent infection in TCCSUP and EJ28 cell lines. Comparisons of persistent TCCSUPPi and control TCCSUP (TCCSUPPi vs. TCCCSUP) revealed 63 significantly DEGs as shown in the Volcano plot (Supplementary Fig. 4A), with 25 upregulated (log2 fold-change ≥ 0) and 38 downregulated genes (log2 fold-change ≤ 0). Genes found towards the top of the plot are highly statistically significance and those found in the extreme right or left are strongly up and down-regulated respectively. When comparing persistent EJ28Pi and control EJ28, 134 DEGs were identified as shown in the Volcano plot (Supplementary Fig. 4B) with 55 upregulated (log2 fold-change ≥ 0) and 79 downregulated genes (log2 fold-change ≤ 0). EJ28Pi versus TCCSUPPi successfully identified 786 DEGs, including 693 upregulated (log2 fold-change ≥ 0) and 93 downregulated genes (log2 fold-change ≤ 0) as illustrated in the Volcano plot (Supplementary Fig. 4C).

The number of downregulated genes was found to be higher than upregulated genes when comparing persistently infected cells with their control (Fig. 2A). In contrast, the comparison between EJ28Pi and TCCSUPPi cells showed that the number of upregulated genes were higher than downregulated genes (Fig. 2A). Furthermore, EJ28Pi had the highest number (78) of DEGs while TCCSUPPi had the lowest number (7) of DEGs. Alteration in transcripts abundance estimate and the number of uniquely expressed genes were visualized in Venn diagram (Fig. 2B). Profiles of the top 20 DEGs among EJ28Pi vs. TCCSUPPi, TCCSUPPi vs. TCCSUP, and EJ28Pi vs. EJ28 libraries were represented in hierarchical clustering based on FPKM log ratio data (FDR <0.05) as illustrated in Fig. 2C, Fig. 2D and Fig. 2E. The red colour represents high expression and orange colour represents lower expression.

**Fig. 2.**
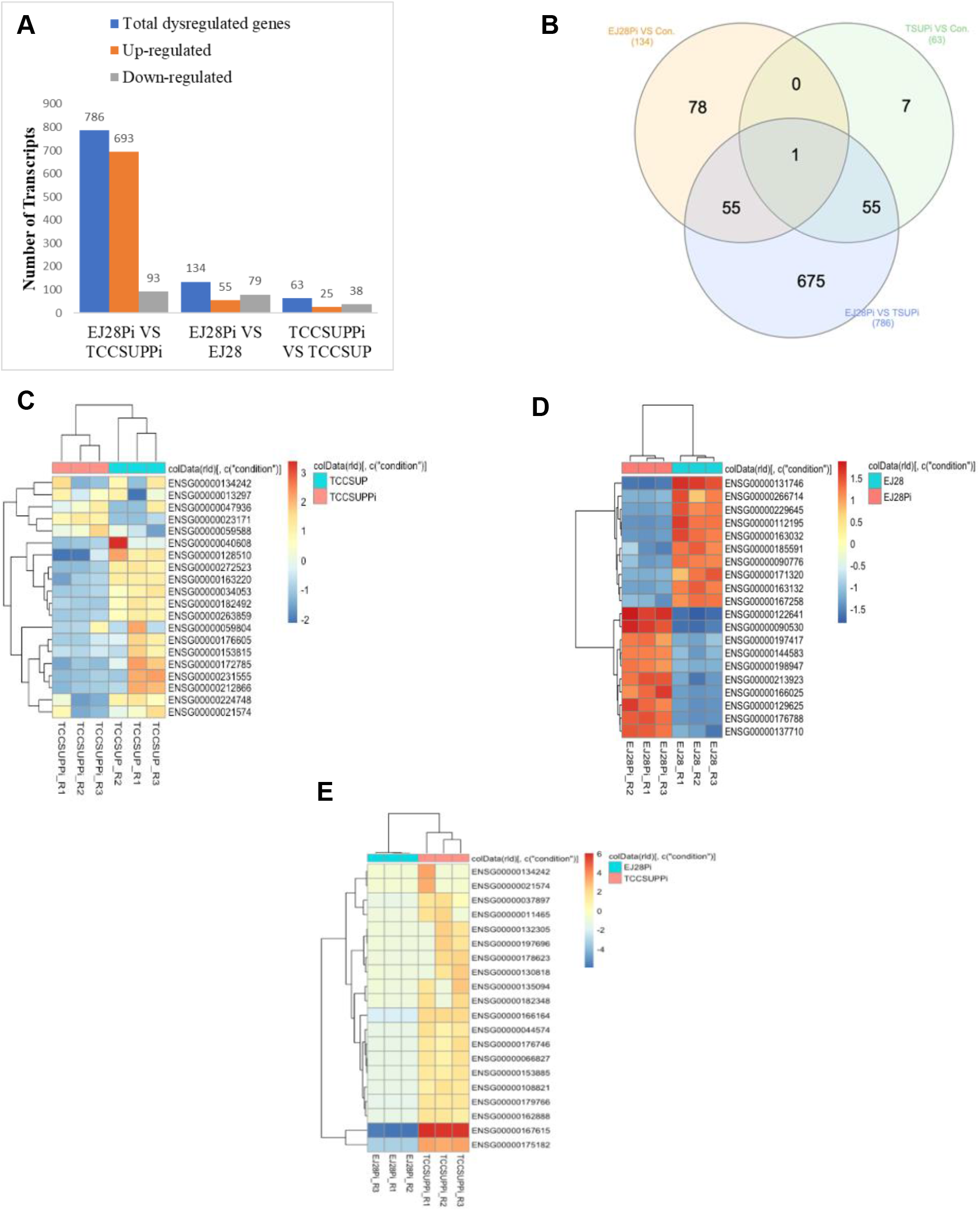
Summary of DEGs among persistently infected and uninfected cells. A) Changes in gene expression profile among EJ28 and TCCSUP persistently infected libraries. Total number of dysregulated genes in the global transcriptome, number of up-regulated and down-regulated genes between EJ28Pi versus TCCSUPPi, EJ28Pi versus EJ28 and, TCCSUPPi versus TCCSUP are summarized. B) Venn diagram showing differentially expressed genes that are among the three libraries of EJ28Pi and TCCSUPPi. EJ28Pi VS Con refers to comparison between EJ28Pi and EJ28 libraries, TSUPi VS Con is comparison between TCCSUPPi and TCCSUP libraries. EJ28Pi VS TSUPi refers to comparison between EJ28Pi and TCCSUPPi libraries. C) Hierarchical clustering of top 20 differentially expressed genes in TCCSUPPi versus TCCSUP libraries. D) Hierarchical clustering of top 20 differentially expressed genes in EJ28Pi versus EJ28 libraries. E) Hierarchical clustering of top 20 differentially expressed genes among the EJ28Pi versus TCCSUPPi libraries. Profiles of the top 20 DEGs among all the six libraries was based on log ratio FPKM data (FDR <0.05) and absolute value of the log2 ratio >2). The horizontal bar above the dendrogram represents untreated (turquoise blue) and treated (dark peach) samples. Level of gene expression was indicated by a range between red (high expression) and blue (low expression).

The number of genes that was altered due to persistent NDV infection in EJ28Pi cells were found to be greater by two-fold than those observed in TCCSUPPi cells with only 0.51% number of co-regulated gene (*UPK3BL1*). Although *UPK3BL1* gene is found to be co-expressively regulated in both the NDV persistently infected cell lines, it was observed to be upregulated in TCCSUPPi but downregulated in EJ28Pi. Thus, the finding suggests that there is a great difference in transcripts expression between NDV persistent infection in TCCSUPPi and EJ28Pi cells. The top 4 genes found to be upregulated in TCCSUPPi are *ACER3*, *THBS1*, *UPK3BL1* and *GRAMD1B*. Those that were upregulated in EJ28Pi are *INHBA*, *P3H2*, *BASP1*, and *AMOTL1*. On the other hand, *APBA2*, *HSPA1B*, *CBWD1* and *BGN* genes were identified to be downregulated in TCCSUPPi and *TNS4*, *TREML2*, *VSNL1*, *MSX1* genes were downregulated in EJ28Pi.

### Functional annotation of the DEGs

To further investigate the biological significance of the identified putative differentially expressed genes (DEGs) in the two-way comparisons of the libraries from EJ28Pi vs TCCSUPPi, TCCSUPPi vs TCCSUP, and EJ28Pi vs EJ28, statistically significant differentially expressed transcripts with *p* <0.05 were used to test against the background set of all genes with GO annotations in metascape[33]. A total of 55 GO terms were discovered in EJ28Pi vs TCCSUPPi and grouped into three ontological categories; biological process (BP), cellular component (CC), and molecular function (MF). Based on this GO terms clustering, the GO terms distribution was equal in biological process (20 GO terms) and molecular function (20 GO terms) and finally, the cellular component (15 GO terms). The significantly enriched GO terms for biological processes in EJ28Pi vs TCCSUPPi were cell cycle G1/S phase transition (GO: 0044843, *p* = 0.0001) and G1/S transition of mitotic cell cycle (GO: 0000082, *p* = 0.0001), while those for cellular component were transport vesicle (GO: 0030133, *p* = 0.0001) and membrane coat (GO: 0030117, *p* = 0.0001), and those for molecular function were intracellular calcium activated chloride channel activity (GO: 0005229, *p* = 0.0001) and phosphatidylserine binding (GO: 0001786, *p* = 0.0001) (Fig. 3; Supplementary Table 2).

**Fig. 3.**
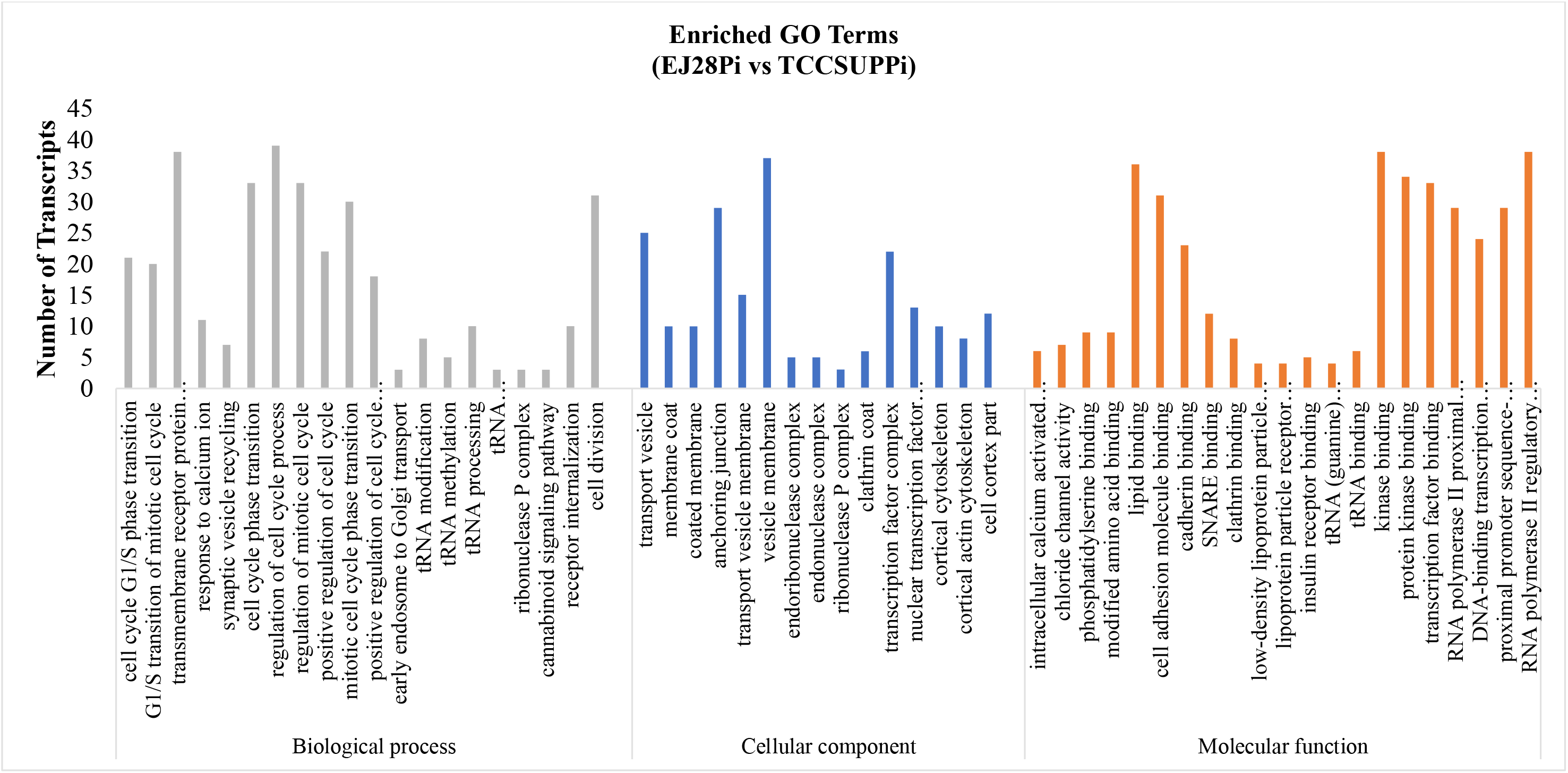
Gene ontology (GO) functional enrichment analysis of DEGs in EJ28Pi relative to TCCSUPPi. The function of genes identified cover three main levels of gene ontology categories: biological process (green), cellular component(blue), and molecular function (red). The y-axis represents the number of transcripts and x-axis represents the ontology categories.

A total of 22 GO terms were respectively clustered into biological process (12 GO terms), cellular component (6 GO terms) and molecular function (4 GO terms) in TCCSUPPi vs TCCSUP. The GO terms found to be significantly enriched in the persistent TCCSUPPi cells for biological processes were response to unfolded protein (GO: 0006986, *p* = 0.0001) and response to topologically incorrect protein (GO: 0035966, *p* = 0.0001), while those for cellular component were collagen-containing extracellular matrix (GO: 0062023, *p* = 0.0001) and extracellular matrix (GO: 0031012, *p* = 0.0001) and those found in molecular function were calcium ion binding (GO: 0005509, *p* = 0.0001) and DNA-binding transcription repressor activity, RNA polymerase II-specific (GO: 0001227, *p* = 0.0001) (Fig.4; Supplementary Table 3).

**Fig. 4.**
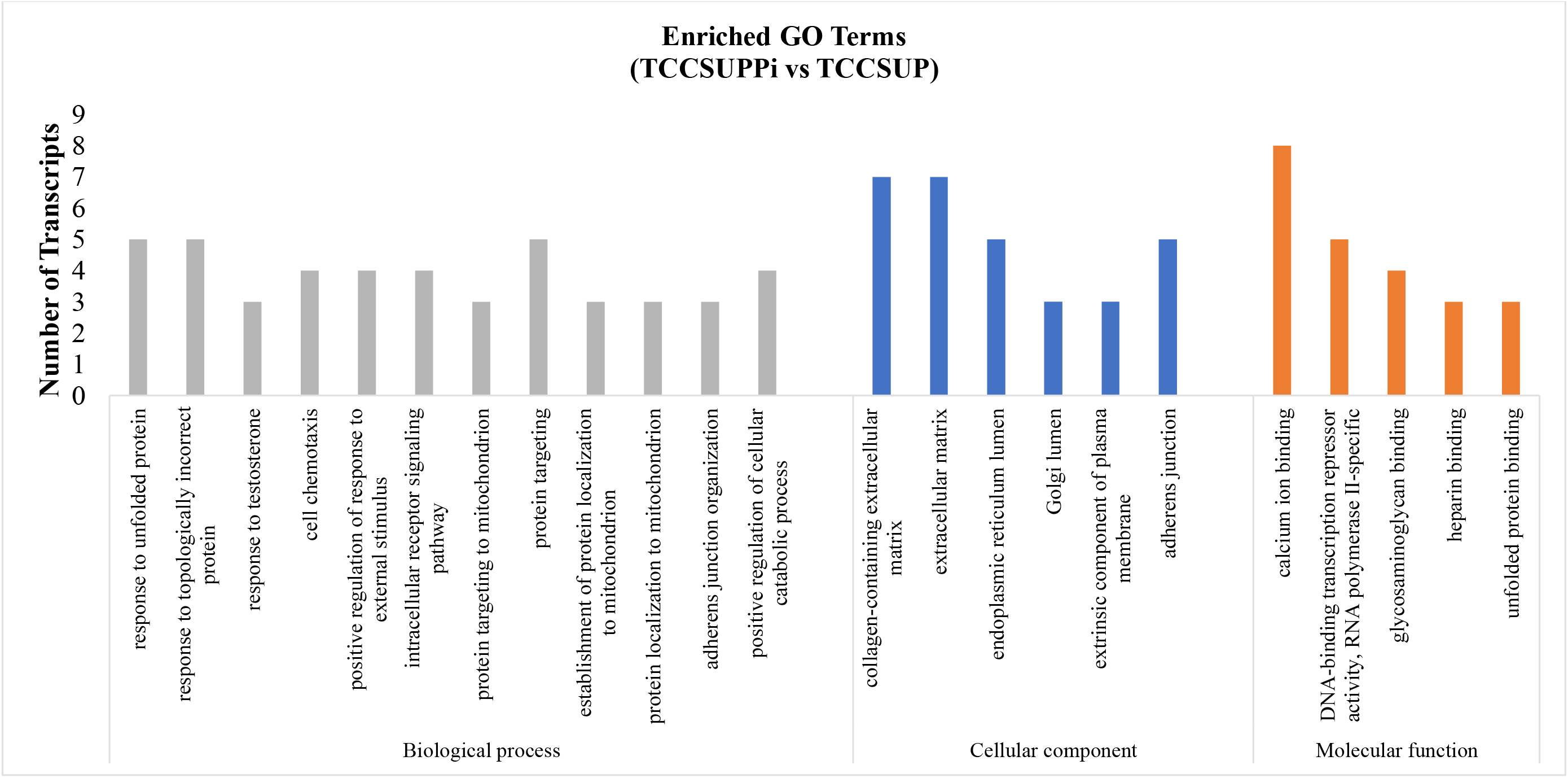
Gene ontology (GO) functional enrichment analysis of DEGs in TCCSUPPi relative to control. The function of genes identified cover three main levels of gene ontology categories: biological process (green), cellular component(blue), and molecular function (red). The y-axis represents the number of transcripts and x-axis represents the ontology categories.

While for EJ28Pi vs EJ28, a total of 44 GO terms were discovered and also respectively clustered into biological process (20 GO terms), cellular component (4 GO terms) and molecular function (19 GO terms). The significantly enriched GO terms found in persistent EJ28Pi cells for biological process were negative regulation of cell proliferation (GO: 0008285, *p* = 1.04E-03) and head development (GO: 0060322, *p* = 1.01E-03), whereas cellular component were membrane region (GO: 0098589, *p* = 0.0001) and membrane raft (GO: 0098589, *p* = 0.0001) and those in molecular function were intracellular calcium activated chloride channel activity (GO: 0005229, *p* = 1.01E-03) and phosphatidylserine binding (GO: 0001786, *p* = 1.03E-03) (Fig.5; Supplementary Table 4). Overall, the results demonstrate that changes in biological processes may play a vital role in persistently infected EJ28 bladder cancer cell lines.

**Fig. 5.**
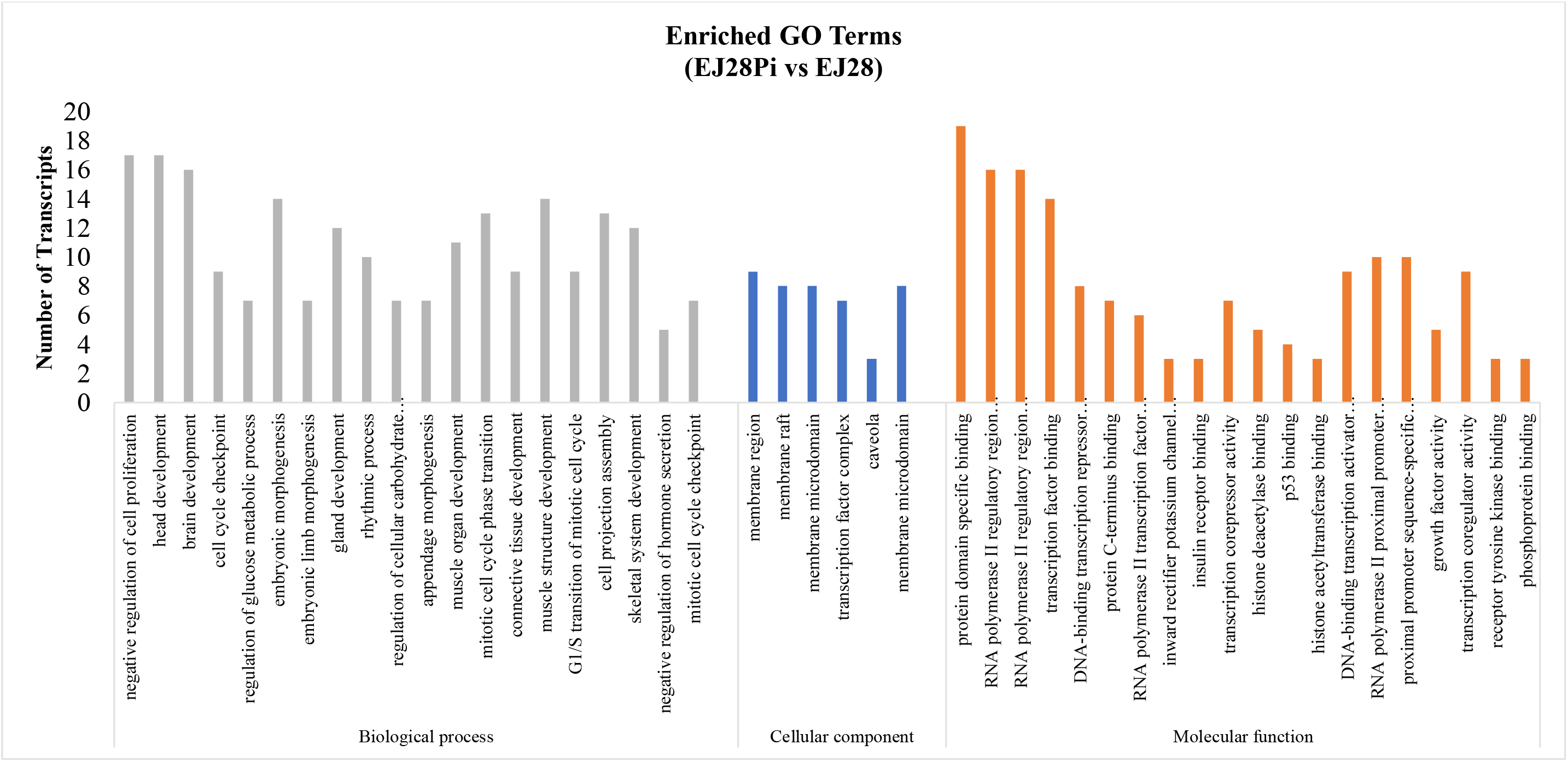
Gene ontology (GO) functional enrichment analysis of DEGs in EJ28Pi relative to control. The function of genes identified cover three main levels of gene ontology categories: biological process (green), cellular component(blue), and molecular function (red). The y-axis represents the number of transcripts and x-axis represents the ontology categories.

### Hallmark pathways enrichment analysis

To understand more about the functions of differentially expressed transcripts in response to NDV persistent infection in TCCSUP and EJ28 bladder cancer cell lines, DEGs were mapped against the most widely used gene set database for performing comprehensive gene set enrichment analysis [34], Molecular Signatures Database (MSigDB) and compared with the whole transcriptome background using Fast Gene Set Enrichment Analysis (FGSEA) package in R [35]. The analysis identified a total of 54 dysregulated pathways due to persistent infection in TCCSUPPi and EJ28Pi cell lines with 31 upregulated and 23 downregulated pathways (Supplementary Fig. 5A). From each persistently infected cell analysis, 15 pathways were found to be affected in TCCSUPPi cells with 9 upregulated and 6 downregulated pathways (Supplementary Fig. 5B while 39 pathways were dysregulated in EJ28Pi with 22 upregulated and 17 downregulated pathways (Supplementary Fig. 5C). In both persistently infected cell lines, many dysregulated pathways were upregulated than downregulated, suggesting an increase level of cellular and physiological response. Twelve (12) hallmark pathways from GSEA common to either TCCSUPPi or EJ28Pi were enriched as demonstrated in TCCSUPPi vs EJ28Pi combined pathways analysis (Table 1). These pathways are allograft rejection, hypoxia, epithelial mesenchymal transition, p13k-Akt_MTOR signaling, heme metabolism, complement, coagulation, estrogen response late, apoptosis, angiogenesis, xenobiotic metabolism and MTORC1 signaling (Table 1). However, the commonly shared pathways such as hypoxia, complement, estrogen response late and MTORC1 signaling pathways were found downregulated in TCCSUPPi cells but upregulated in EJ28Pi cells which suggest that bladder cancer cell lines response to NDV persistent infection differently.

**Table 1:** Twelve (12) hallmark pathways from GSEA commonly shared among TCCSUPPi and EJ28Pi.

| S/N. |  | Pathway shared by TCCSUPPi & EJ28Pi | Total | DEGs in TCCSUPPi | DEGs in EJ28Pi |
| --- | --- | --- | --- | --- | --- |
| 1 | Upregulated pathways | Allograft rejection | 200 | *F2* | *ETS1, INHBA* |
| 2 |  | Hypoxia | 200 | *BGN* | *ETS1, BHLHE40* |
| 3 |  | Epithelial mesenchymal transition | 200 | *CDH2, THBS1* | *INHBA, BASP1* |
| 4 |  | p13k-Akt_MTOR signalling | 105 | *CALR* | *SFN, RAC1, PTPN11* |
| 5 |  | Heme metabolism | 200 | *BACH1* | *MARK3* |
| 6 |  | Complement | 200↓ | *S100A9* | *ZEB1, OLR1* |
| 7 |  | Coagulation |  | *THBS1, F2* | *RAC1, OLR1* |
| 8 |  | Estrogen response late | 200↓ | *S100A9* | *SFN, HOMER2, EMP2, FARP1* |
| 9 | Downregulated pathways | Apoptosis | 161 | *BGN* | *CCND1, FASLG, WEE1* |
| 10 |  | Angiogenesis | 36 | *LRPAP1* | *MSX1, PDGFA, OLR1* |
| 11 |  | Xenobiotic metabolism | 200 | *HES6* | *SLC1A5* |
| 12 |  | MTORC1 signalling | 200↑ | *HSPA4, CALR* | *SLC1A5* |

**Note:** The arrow up (↑) indicates that the pathway is upregulated in TCCSUPPi and the arrow down (↓) indicates that the pathway is downregulated in TCCSUPPi.

The major upregulated pathways in TCCSUPPi were heme metabolism, TGF-beta signaling and spermatogenesis (Supplementary Fig. 5B), while the downregulated pathways were estrogen late response, compliment and hypoxia. On the other hand, allograft rejection, KRAS signaling and interferon gamma response were observed to be upregulated in EJ28Pi while MTORC1 signaling, xenobiotic metabolism and IL2 STAT5 signaling were discovered to be downregulated (Supplementary Fig. 5C). Interestingly, apoptosis pathway was commonly observed to be downregulated in both persistently infected TCCSUPPi and EJ28Pi cells (Supplementary Fig. 5A) and that could be the reason why NDV persistently infected cancer cell lines maintains their normal growth and proliferation. It is well known that NDV induces apoptosis and kills cancer cells through activation of apoptotic signaling pathways thus it is an exciting discovery that this pathway is downregulated in the established persistently infected cell lines.

### Calcium ion binding genes in TCCSUPPi

Two classical and important cadherin superfamily genes were significantly expressed in NDV persistently infected TCCSUPPi cells (FDR <0.05). These genes are cadherin 2 (*CDH2*) and cadherin 5 (*CDH5*) with *CDH2* upregulated at 0.92 fold and *CDH5* downregulated at −2.21 fold. Both cadherin 2 (*CDH2*) and cadherin 5 (*CDH5*) are protein-coding type of genes and are calcium-dependent cell-cell adhesion molecules that encode five extracellular cadherin repeats known as transmembrane region. They allow cells to make homophilic interactions and play a vital role in endothelial adherens junction assembly and maintenance, making the cells more connected to the actin cytoskeleton through formation of alpha-catenin [36]. Moreover, these key genes have been reported to be involved in tissue morphogenesis[37, 38] and their misexpression is implicated in human cancer[39–41]. Thus, it is likely that upregulation of *CDH2* may be associated with increased interconnection between the persistent cells leading to the maintenance of cellular structure, cell mobility and intercellular transport[42], and downregulation of *CDH5* may lead to reduced calcium ion binding and intercellular junctions disorganization in persistent TCCSUPPi.

### Repression of stress proteins and transcriptional genes in TCCSUPPi

Genes involved in heat shock proteins and homeobox subfamilies were revealed to be downregulated in NDV persistent TCCSUPPi cells. The downregulated genes found for heat shock includes *HSPA1B*, *HSPA4* and *HSPB7,* and those of homeobox family were *HOXA13* and *HOXB13* (Supplementary Table 5). Expression of heat shock protein family A (Hsp70) member 1B (*HSPA1B*) and, heat shock protein family A (Hsp70) member 4 (*HSPA4*) as well as heat shock protein family B (small) member 7 (*HSPB7*) were reported to be frequently overexpressed in bladder cancer cell lines[43, 44]. This is due to the remarkable differences in bladder cancer expression compared with normal urothelium[45], leading to an intense investigation and suggestion for biomarkers of bladder cancer treatment response and prognosis[46]. Thus, since the heat shock proteins were found to be significantly downregulated (FDR <0.05) in TCCSUPPi cells, it can be said that NDV may be utilized to specifically target those family genes to enhance therapeutic strategies for patients with bladder cancer.

Additionally, homeobox transcription factor gene family such as *HOXA13* and *HOXB13* were also shown to be downregulated in persistent TCCSUPPi cells. These genes are members of homeobox family which encode the homeodomain proteins that play an important role in Mullerian duct development, gut and urogenital tract[47, 48]. Repression or expression of HOX genes have both been reported to be linked with cancer where they act as either tumour suppressor or proto-oncogenes[49, 50]. Mutations of *HOXA13* gene have been reported to be associated with abnormalities in mice and a serious uterine development effect in human. *HOXA13* has been described to have the same regional level of expression in both adult and embryo[47, 51]. Both *HOXA13* and *HOXB13* genes act to provide protein that bind to a specific region of DNA and co-regulate functional activity of other genes[52]. The proteins produced by these set of genes are called transcription factors that are often referred to as homeobox protein family. Many RNA viruses (such as micro RNA virus, influenza, vesicular stomatitis virus, rhabdovirus etc.) were reported to inhibit host cell gene expression [53, 54] with NDV inhibiting transcription inhibitor genes[55]. Based on this repression of transcriptional and stress protein genes in TCCSUPPi cells, it can be hypothesized that NDV requires reduction of mRNA output of cells for its effective replication in cancer cells.

### Novel apoptosis, cell cycle and immune-related genes in EJ28Pi

It is established that infection of cancer cells with NDV trigger an innate immune response that leads to apoptosis and cell death, making NDV an efficient oncolytic mediator. In this study, four novel apoptosis related genes, including etoposide-induced gene 24 (*EI24*), epithelial membrane protein 2 (*EMP2*), tumour protein p53 (*TP53*), and prostaglandin E synthase (*PTGES*) were revealed to be significantly downregulated in EJ28Pi cells (FDR <0.05) (Supplementary Table 6). *EI24* gene is known to suppress cell growth by inducing apoptosis through activation of caspase 9 and mitochondrial pathways[56, 57]. *EMP2* gene encodes tetraspan protein of PMP22/EMP family and functioning to signaled cell death and apoptosis[58], overexpression of this gene has resulted in cancer progressions in various types of tissues[59, 60]. *TP53* gene is activated in response to cellular stress to help in regulating the expression of the target genes by inducing cell cycle arrest, DNA repair and apoptosis[61] while *PTGES* is reported to induce proinflammatory cytokine interleukin 1 beta (*IL1B*) and involve in inducing apoptosis via *TP53* gene activation.

Furthermore, three novel apoptosis and immune-related genes such as fas ligand (*FASLG*), sp1 transcription factor (*SP1*), signal transducer and activator of transcription 6 (*STAT6*) were also significantly downregulated (FDR <0.05) in EJ28Pi cells (Supplementary Table 6), suggesting that the downregulation of these apoptosis, and immune-related genes are the possible mechanism employed by EJ28 bladder cancer cells in reducing the negative impact of apoptosis damage usually induced by NDV infection, making it survive NDV infection as demonstrated by the upregulation of genes related to cell cycle, cell growth and cell survival such as MAX network transcriptional repressor (*MNT*), NDRG family member 4 (*NDRG4*), zinc finger protein 346 (*ZNF346*), fibroblast growth factor 14 (*FGF14*), and fibroblast growth factor 9 (*FGF9*) (Supplementary Table 6).

### Solute carrier and potassium voltage-gated channel genes in EJ28Pi

Three solute carrier superfamily genes (*SLC*) including solute carrier family 1 member 5 (*SLC1A5*), solute carrier family 43 member 3 (*SLC43A3*), solute carrier family 48 member 1 (*SLC48A1*) were observed to be significantly downregulated in EJ28Pi cells (FDR <0.05) (Supplementary Table 7). These solute carriers genes (*SLC*) encode multiple transmembrane transporters mostly located in the cellular membrane and are responsible for transportation of glucose, ions amino acids, minerals, vitamin and nucleotides across cell membrane [62, 63]. The results suggest that the NDV persistent infection on EJ28 cells suppresses the transmembrane receptors by downregulating these set of SLC genes thus affecting the oncolytic activity of NDV might have been reduced or the cells have eliminated the virus from its cytoplasm, making it less energy or nutrient demanding. In addition, three members of the inward rectifier-type potassium channel family such as potassium voltage-gated channel subfamily J member 10 (*KCNJ10*), potassium voltage-gated channel subfamily J member 12 (*KCNJ12*), potassium voltage-gated channel subfamily J member 5 (*KCNJ5*) were significantly downregulated in persistent EJ28Pi cells (FDR <0.05) (Supplementary Table 7), demonstrating that the EJ28 bladder cancer cell lines do not require inflow or potassium buffering cation in the cells to develop NDV persistent infection.

### Validation of RNA-Seq data by qPCR

To validate the expression of RNA-Seq data obtained from the persistent TCCSUPPi and EJ28Pi cells, three genes with annotations from statistical analysis of RNA-Seq were randomly selected from the TCCSUPPi RNA-Seq data and two genes from EJ28Pi RNA-Seq data. The selected genes are *HOXA13*, *HOXB13* and *OTX1*. RT-qPCR analysis showed similar expression patterns to that observed in RNA-Seq analysis of TCCSUPPi cells (Fig. 6A). We found *HOXA13*, *HOXB13* and *OTX1* to be significantly downregulated in TCCSUPPi cells relative to TCCSUP cells with fold change values of −40.38, −2.52 and −3.01, respectively (*p*<0.05). On the other hand, *HOXB13* and *HOXA13* genes were found to be significantly upregulated in EJ28Pi cells (3.25-fold change and 2.805-fold change) compared with EJ28 cells (*p*<0.05) (Fig. 6B). These results demonstrated that the change in gene expression deduced by RNA-Seq reflected the actual transcriptome differences between persistently infected bladder cancer cell lines (TCCSUPPi and EJ28Pi) and that of the uninfected control cells (TCCSUP and EJ28). Thus, confirming that the NGS data was accurately quantified and valid for inferring biological significance.

**Fig. 6.**
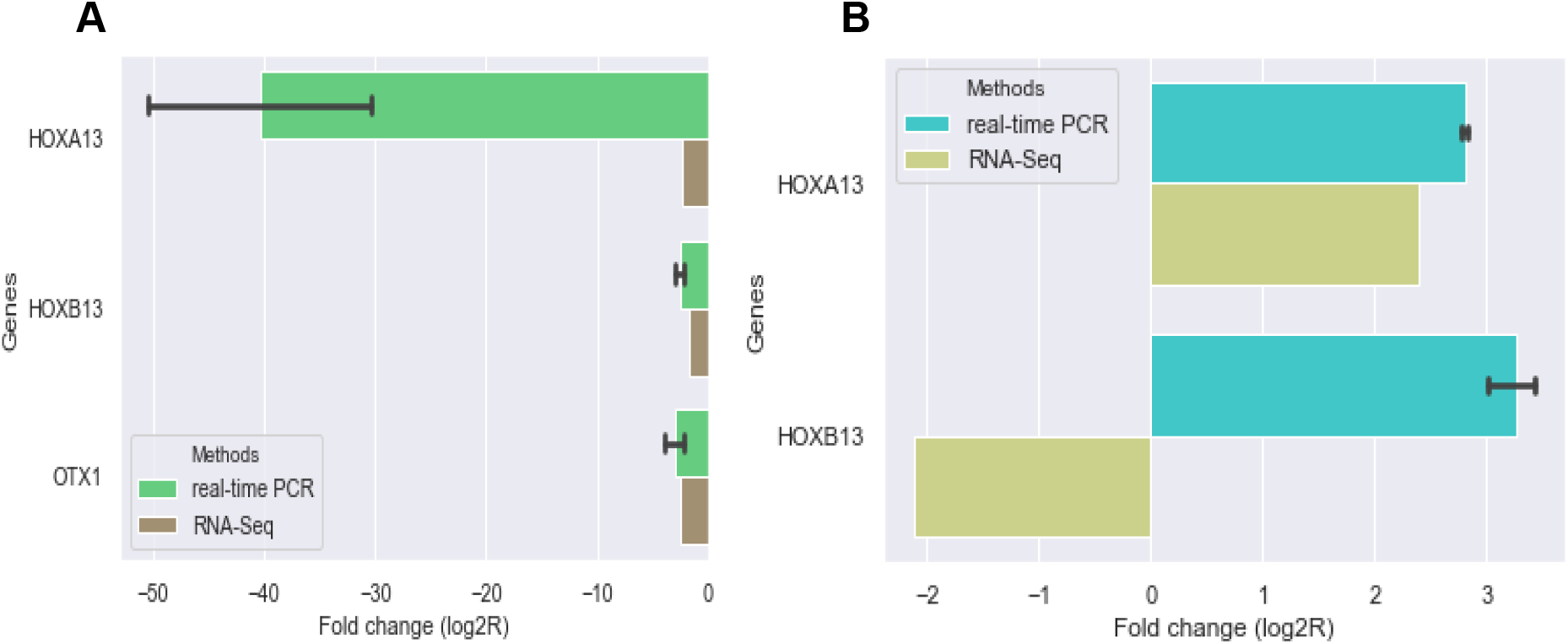
Validation of RNA-seq data of HOXA13, HOXB13, and OTX1 genes by RT-qPCR analysis. A) Comparison of expression level detected by RNA-seq and RT-qPCR in TCCSUPPi compared to TCCSUP cells. B) Comparison of expression level detected by RNA-seq and RT-qPCR in EJ28Pi compared to EJ28 cells. The x-axis shows the annotations of the selected genes while the y-axis shows the normalized expression levels of the transcripts.

## Discussion

To better understand the mechanisms by which bladder cancer cell lines acquire NDV persistent infection, we employed RNA sequencing to profile the cellular transcriptome of the NDV-persistently infected TCCSUPPi and EJ28Pi cell lines compared to their uninfected controls. Our results indicated that there were considerable differences in the level of gene expressions between persistently infected bladder cancer cell lines (TCCSUPPi and EJ28Pi) and their respective uninfected controls (TCCSUP and EJ28), and that several genes were uniquely differentially expressed in the NDV-persistently infected cells. We were able to discover more than (>190) differentially expressed genes (DEGs) associated with NDV persistent infection when compared with previous study using microarray where there were approximately more than (>300) DEGs identified [64]. This could be due to the use of different mRNA quantification techniques and variation in the use of NDV dose. In addition, the high number of gene expression identified in persistent EJ28Pi relative to those identified in persistent TCCSUPPi cells (Figure 2A & B) could be attributed to the fact that each bladder cancer cell line contains a heterogenous subset of cells that respond differently to NDV-mediated oncolysis.

Our findings on persistently infected TCCSUPPi corroborate with previous report [64] where some genes such as *CDH2* and *CDH5* cadherin superfamily are involves in calcium binding function. These key genes have been reported to be involved in tissue morphogenesis[37, 38] and their dysregulated expression have been implicated in human cancer[39–41]. *CDH2* is often referred to as cadherin switching encoding N-cadherin and is known to regulate many biological processes[65, 66]. Our results reveal that this gene is upregulated in persistent TCCSUPPi cells and is consistent with the previous findings that reported *CDH2* gene is to be commonly upregulated in bladder cancer[67]. Moreover, *CDH2* gene is found to play a significant role in the transition between epithelial and mesenchymal cells (EMT) [68]. On the other hand, *CDH5* gene plays a role in the morphology of a vascular muscle and stability, and is also reported to be overexpressed in aggressive human cutaneous melanoma cells as compared to nonaggressive melanoma [69]. Herein, we found that this gene was significantly downregulated in persistent TCCSUPPi cells making the cells less aggressive because repression of *CDH5* gene is associated with lack of vascular-like formation networks [70], implying that dysregulated expression of this gene in persistent TCCSUPPi cells may disrupt the formation of vasculogenic networks as observed in melanoma.

Other important genes that were implicated in TCCSUPPi are stress proteins such as *HSPA1B*, *HSPA4* and *HSPB7* as well as transcription factor genes comprising of *HOXA13* and *HOXB13,* respectively. Heat shock or stress proteins are expressed in normal cells however their expression are upregulated when triggered by cellular stressors such as cytotoxic agents and hypoxia[45]. These set of genes are overexpressed in various types of cancer including bladder cancer cells. Their varied expression in bladder cancers compared to normal bladder tissue has led to the investigation as being a cancer biomarker. Contrary to this, we observed a significant repression of these heat shock family proteins (*HSPA1B*, *HSPA4* and *HSPB7*) in the established NDV persistent TCCSUPPi cell line, demonstrating that NDV has potential application for bladder cancer treatment strategies by targeting these class of heat shock proteins and may serve synergistic effect with some of the current therapeutic measures of bladder cancers such as chemotherapy and radiation therapy. Moreover, *HOXA13* and *HOXB13* homeobox transcription genes were also repressed in NDV persistent TCCSUPPi cells. Repression or overexpression of HOX genes have both been reported to be linked with cancer where they act as either tumour suppressor or proto-oncogenes[49, 50]. *HOXA13* and *HOXB13* genes provide protein that bind to a specific region of DNA and co-regulate the functional activity of other genes in cells [52]. Therefore, we hypothesize that repression of transcriptional and stress proteins genes in TCCSUPPi cells may be required for NDV to effectively replicate and continue its cycle within the host cells. Further study needs to be carried out to unravel the relationship of NDV and repression of these genes in bladder cancer cells.

On the other hand, the DEGs found to be implicated in EJ28Pi were genes associated with apoptosis such as *EI24*, *EMP2*, *TP53* and *PTGES*, cell cycle and growth such as *MNT*, *NDRG4*, *ZNF346*, *FGF14* and *FGF9*, immune related genes that included *FASLG*, *SP1* and *STAT6*, as well as solute carrier family *SLC1A5*, *SCL43A3*, and *SLC48A1* and potassium voltage-gated channel genes comprising of *KCNJ10*, *KCNJ12* and *KCNJ5*. NDV is reported to mediate its oncolytic activity by inducing apoptosis in many types of cancer cells[18, 71–73] and induction of apoptosis in cancer may lead to upregulation of various kind of apoptosis-related genes[74, 75] and apoptosis-related pathways[76]. Contrary to our findings, we observed that apoptosis-related genes were significantly downregulated in NDV persistent EJ28Pi cells. Moreover, *FASLG*, *SP1* and *STAT6* immune-related genes were significantly repressed in persistent EJ28Pi cells and this may likely be the possible mechanism employed by EJ28 bladder cancer cells in evading apoptotic damage induced by NDV infection. This allows the EJ28 cells to survive NDV-mediated oncolysis and develop persistent infection as demonstrated by the upregulation of genes related to cell cycle, cell growth and cell survival such as *MNT*, *NDRG4*, *ZNF346*, *FGF14* and *FGF9*.

On the basis of functional annotation analysis of the DEGs identified in persistent TCCSUPPi cells, we found that the significantly enriched GO terms were response to unfolded protein and topologically incorrect protein for biological processes, collagen-containing extracellular matrix and endoplasmic reticulum for cellular component, calcium ion biding and DNA-binding transcription repressor activity, RNA polymerase II specific for molecular function. The results of GO term for biological process and cellular component in our findings is consistent with the previous studies which reported that NDV induces autophagy to favour its replication in cell via activation of cellular stress that lead to interference with endoplasmic reticulum (ER) function resulting in unfolding of proteins in ER, a term known as ER stress[77, 78]. Unfolded protein response is a process by which ER employs certain mechanism to prevent cell death[79]. Recent findings have demonstrated that this event may be associated with response to autophagy which plays a vital role in response to stress stimuli like oncolytic viruses such as NDV [80, 81].

On the other hand, the identified DEGs in persistent EJ28Pi cells were significantly enriched with the GO terms such as negative regulation of cell proliferation and cell cycle check point for biological processes, membrane region and membrane raft for cellular component, protein domain specific binding and RNA polymerase II regulatory region sequence-specific DNA binding for molecular function. Viruses either directly damage the cell’s DNA or disrupt basic mechanism of cell division such as the segregation of chromosome and DNA replication[82]. This usually results in the activation of cell cycle checkpoints, a mechanism implemented by cells to repair damage or commit suicide when the cell’s damage fails to repair. In the present study, we found that cell cycle checkpoint is activated as part of the biological event occurring due to NDV persistent infection in EJ28 bladder cancer cell line. Studies have localized the presence of viral structural proteins in both the membrane region and membrane raft of the host cell and demonstrated their contribution for virus entry, assembly and budding[83]. In NDV persistent EJ28Pi cells, the cellular involvement of membrane raft is obvious, suggesting that NDV may be using the membrane raft component of the host cell in facilitating its replication and infectious cycle processes to persistently infect the EJ28 cells through protein domain specific binding and RNA polymerase II regulatory sequence-specific DNA binding.

It is interesting to note that the enriched GO terms in persistent TCCSUPPi cells were entirely different from those found in persistent EJ28Pi cells. This suggests that NDV persistent infection in different bladder cancer cell lines involve different sets of biological processes, cellular component and molecular function for each individual cell line. However, the exact mechanism for this difference is unclear and may need further investigation.

In terms of the disrupted molecular pathways that were enriched due to NDV persistent infection in these two bladder cancer cell lines, 15 pathways were dysregulated in persistent TCCSUPPi cells with 9 upregulated and 6 downregulated pathways. The top three most enriched upregulated pathways in this cell were heme metabolism, TGF-beta signaling and spermatogenesis while the downregulated pathways were estrogen response late, complement, and hypoxia. These annotated pathways provided valuable insight in revealing the molecular cascades interaction for TCCSUP bladder cancer cell line to develop NDV persistent infection. One important pathway involved in NDV persistent infection in TCCSUP cells is TGF-beta signaling, which is an essential pathway that regulates many cellular processes such as cell growth, proliferation, differentiation and migration[84], and dysregulation of this pathway contributes to various types of pathologies that includes cancer. It is reported that enrichment of TGF-beta signaling is a result of an induced action of epithelial-mesenchymal transition (EMT) that facilitates tumour migration and invasion[85]. Moreover, TGF-beta signaling is associated with release of mitogenic growth factor that stimulate proliferation and survival of cancer which correlates with cancer angiogenesis, and metastasis[85]. Therefore, data from TCCSUPPi provide new ideas and strategies employed by TCCSUP cells in acquiring NDV persistent infection.

Apart from that, 39 pathways were implicated in persistent EJ28Pi cells with 22 upregulated and 17 downregulated pathways. Among these dysregulated enriched pathways were allograft rejection, KRAS signaling up and interferon gamma response while those that were downregulated are MTORC1 signaling, xenobiotic metabolism, and IL2 STAT5 signaling. Indeed, NDV is known as a potent stimulator of type I interferon but defect in interferon signaling enhances NDV replicative ability in cancer cells and this has been the basis for its selective replication in cancer. Evidence of proinflammatory cytokine particularly type I IFN production upon NDV infection in both normal and cancer cells has been document [86–89]. However, differences in the level of proinflammatory cytokine between cancer cells and normal tissue as well as their role in NDV-mediated oncolysis is not clearly understood. Taken together, twelve (12) commonly identified dysregulated pathways shared by both TCCSUPPi and EJ28Pi are angiogenesis, xenobiotic metabolism, allograft rejection, hypoxia, epithelial mesenchymal transition, p13k-Akt_MTOR signaling, heme metabolism, complement, coagulation, estrogen response late, apoptosis, and MTORC1 signaling. However, among the commonly shared pathways like; hypoxia, complement, estrogen response late and MTORC1 signaling were downregulated in the persistent TCCSUPPi cells and upregulated in EJ28Pi cells, affirming that bladder cancer cell lines response to NDV persistent infection differently. Therefore, this implies that the important pathways influenced by NDV persistent infection in TCCSUP and EJ28 bladder cancer cell lines are heme metabolism and TGF beta signaling and KRAS signaling up and interferon gamma response, suggesting that the genes in these pathways contribute significantly to the development of NDV persistent infection in bladder cancer cell lines.

To further validate the transcriptome data, we randomly selected three DEGs for validation by RT-qPCR in TCCSUPPi and TCCSUP, and two DEGs in EJ28Pi and EJ28 RNA-Seq dataset. Consistent with those of the RNA-Seq datasets, we discovered *HOXA13*, *HOXB13*, *OTX1* were differentially expressed between persistent TCCSUPPi and TCCSUP control cells. Similarly, *HOXB13* was also found to be differentially expressed between persistent EJ28Pi and EJ28 control cells. *HOXA13*, *HOXB13* and *OTX1* genes were downregulated in TCCSUPPi cells and *HOXB13* gene was also downregulated in EJ28Pi cells. HOX genes also known as homologous genes are a subset of homeobox genes family that function in regulating morphogenesis in living cells. Both *HOXA13* and *HOXB13* are gene members of HOX family. Their role in cancer has been widely studied[90–92] and these genes were implicated in bladder cancer progression[93, 94] and poor prognosis[95]. The different expression of *HOXA13*, *HOXB13* and *OTX1* genes between TCCSUPPi and TCCSUP as well as HOXB*13* gene between EJ28Pi and EJ28 suggest that regulatory mechanism of these genes varied between these cell lines. Findings from this study reflected the actual differences in expression profiles computed by RNA-Seq between NDV persistent bladder cancer cell lines and their uninfected control cells.

## Conclusion

We employed RNA sequencing analysis to identify commonly dysregulated transcriptomic profiles and disrupted molecular pathways in NDV persistently infected bladder cancer cell lines. Through this analysis, we were able to identify several important dysregulated genes as well as pathways that seems to be the key players in development of NDV persistent infection. We reported the discovery of 63 and 134 uniquely expressed transcripts that were differentially expressed in TCCSUPPi and EJ28Pi cells, respectively. The profiles of TCCSUPPi cells were characterized by overexpression of calcium ion binding genes (*CH2*), repression of stress proteins such as *HSPA1B*, *HSPA4*, and *HSPB7*, and downregulation of transcriptional factor gene namely; *HOXA13* and *HOXB13*. In EJ28Pi, downregulation of some novel apoptosis genes that includes *EI24*, *EMP2*, *TP53*, and *PTGES*, repression of immune response genes comprising of *FASLG*, *SP1* and *STAT6* as well as overexpression of cell cycle and growth-related genes including *MNT*, *NDRG4*, *ZNF346*, *FGF14* and *FGF9* and downregulation of solute carrier family such as *SLC1A5*, *SLC43A3* and *SLC48A1* with significant repression of potassium voltage-gated channel genes such as *KCNJ10*, *KCNJ12* and *KCNJ5* were more obvious. These genes may have an important role in the development of NDV persistent infection in bladder cancer cells. Based on gene ontology analysis, we understand that the changes in molecular function are more important in the establishment of NDV persistent infection in bladder cancer cells. As for the pathways, we reported the involvement of heme metabolism, TGF-beta signaling, spermatogenesis, response late, complement, hypoxia, allograft rejection, KRAS signaling up, interferon gamma response, MTORC1 signaling, xenobiotic metabolism, and IL2 STAT5 signaling as the pathways identified in this study to have played a crucial role in development of NDV persistent infection in bladder cancer cells.

## Materials and Methods

### Cells and virus

Two human bladder cancer cell line, TCCSUP and HTB-5, were purchased from the European Collection of Authenticated Cell Cultures (ECACC; Salisbury, UK) while one NDV persistently infected human bladder cancer cell line (designated as EJ28Pi) was provided by Dr. Chia Suet Lin. The cell lines were cultured and maintained in MEM-alpha and RPMI 1640 containing 10% FBS (Gibco, Life Technologies) incubated at 37°C supplied with 5% CO_2_. *Mycoplasma* test was carried out on each cell line to ensure the cells are free from *Mycoplasma* contamination prior to the establishment of cells with NDV persistent infection. The wild-type oncolytic Newcastle disease virus (NDV) AF2240 strain was obtained from the Virology Laboratory of the Faculty of Biotechnology and Biomolecular Sciences, Universiti Putra Malaysia (UPM) and was propagated in a specific pathogen free (SPF) embryonated chicken eggs of 9 to 11 days old at 37∘C for 72 hours as previously described [23]. Allantoic fluid were harvested from the eggs and presence of virus was confirmed by haemagglutination assay[96] while virus purification was based on previously established protocol[97]. The resulting virus pellet were stored in −80°C freezer until use.

### Establishment of NDV persistent infection

Persistently infected cells were developed as described previously [23]. In brief, TCCSUP bladder cancer cells was infected with NDV at MOI of 0.1 in a T25 flask. After an hour of incubation at 37°C for viral absorption, cells were washed with 1X PBS and washed again twice with MEM alpha media to inactivate unabsorbed viruses. Cells were replenished with maintenance medium (MM) constituted with 2% FBS (Gibco, Life Technologies) and incubated for a period of 5 days. Majority of the cells were detached after these post infection days. The cells were then washed twice with 1X PBS and complete growth medium was added. The cells were allowed to grow until sub-confluent monolayer was formed. The newly established NDV persistently infected bladder cancer cell line was designated as TCCSUPPi.

### RNA extraction and quality control

Total RNA was extracted for each of the persistently infected cells as well as the cell lines without NDV infection (negative control) using Qiagen RNeasy micro kit (Qiagen, Germany) following the manufacturer’s protocol. Genomic DNA was removed from the RNA samples by DNase Max treatment (Cat no.: 15200-50, MO Bio Inc.) following manufacturer’s instructions. RNA quality and RNA integrity number (RIN) were assessed by spectrophotometer (NanoDrop 2000) and Bioanalyzer 21000 (Agilent Technologies, Santa Clara, CA, USA), respectively. RNA Integrity Number (RIN) were in the range between 8.0 to 9.0 and thus were of good quality. Quantification of RNA quantification was carried out using Qubit® RNA Assay Kit and Qubit® 2.0 Fluorometer (Life Technologies, San Diego, CA, USA).

### Library constructions and RNA sequencing

Paired-end libraries were constructed by the TruSeq Stranded mRNA library preparation kit v2 (Illumina, San Diego, CA) with a total of 3 μg of RNA per sample. Briefly, oligo-dT beads were used to purify polyadenylated RNA containing mRNA molecules. This was followed by fragmentation of the purified mRNA molecules into smaller fragments whereby random primers were used to reverse-transcribed the RNAs into cDNA using SuperScript II Reverse Transcriptase (Invitrogen, part #18064-014). Following the reverse transcription of the RNA, the fragments were 30-end adenylated and ligated to standard Illumina paired-end cluster generation (sequencing adapters) and finally the bar-coded library products were amplified by 12 cycles of PCR. The final amplified cDNA libraries were analyzed by Bioanalyzer 21000 (Agilent Technologies, Santa Clara, CA, USA) and the electropherogram was checked to have a narrow distribution with at least a peak size of approximately 300 bp. The Illumina TruSeq mRNA libraries with an insert size of about 200 nt were clustered equimolar into a 10nM sequencing stock solution and amplified followed by QC check using 2100 Bioanalyzer (Agilent, Santa Clara, CA). A total of 12 libraries from persistently infected cells and control were generated and sequenced using HiSeq2000 Illumina platform following manufacturer’s instruction (Illumina Inc., San Diego).

### RNA-seq data quality check and processing

#### MultiQC analysis

The FASTQ file for the raw sequence data was subjected to quality control checks (QC) using the FastQC software[98] and the verified sequence data has been submitted to NCBI Gene Expression Omnibus (GEO) with accession number GSE140902. The dataset was available at https://www.ncbi.nlm.nih.gov/geo/query/acc.cgi?acc=GSE140902. The results and the log files (output) generated by FastQC were collected and applied to MultiQC to have an overall summary statistics of the data in a single standalone HTML report file while saving the directory of the remaining parsed data for further downstream application[99].

#### Alignment of the sequencing read to a reference

The FASTQ mapping was performed using HISAT2 that is fast and suitable for splice site alignment[100]. In brief, reference human genome file sequence in FASTA format was obtained from Ensembl genome browser[101] and the sequences were used to map each of the paired-end reads to the hg19 human reference genome by using HISAT2. The mapping rate for the samples were above 95% and shows high mapping rate. Mapping rate were summarized by type to gain an insight into which samples were properly mapped and which was not. Reads that mapped to the hg19 human reference genome was used to quantify the transcriptome assembly using StringTie[102].

### Gene expression analysis

Quantification of the transcripts abundance was carried out using Salmon’s quasimapping strategy which counts the number of reads that is mapped to a given transcript[103] calculated in fragments per kilobase of gene model per million mapped (FPKM) of the coding genes in each samples. Differential transcripts analysis (DEGs) was carried out using DESeq2 package[32] via tximport in R environment.

### Functional and pathway enrichment analysis

Functional enrichment analysis of the differentially expressed genes (DEGs) was carried out using metascape (http://metascape.org/gp/index.html), an online platform for comprehensive gene annotation analysis of omics data[33]. Gene ontology (GO) analysis of the DEGs was ranked by enrichment score of 1/log10 *p* value and curated in MS Excel. Hallmark pathways enrichment analysis was performed using FGSEA R-package for fast preranked gene set enrichment analysis (GSEA)[104] and the background was set using Molecular Signatures Databases (MSigDB)[105]. The identified GO terms and hallmark pathways with a corrected *p* <0.05 were considered significantly enriched.

### RT-qPCR analysis of RNA-Seq data

Total RNAs from both TCCSUPPi and EJ28Pi cells were used for RT-qPCR analysis with uninfected TCCSUP and EJ28 as negative controls. A subset of three genes (*OTX1*, *HOXA13* and *HOXB13*) with annotations from RNA-Seq analysis were randomly selected for RT-qPCR analysis. In brief, each RNA sample was converted to cDNA using QuantiTect Reverse Transcriptase kit (Qiagen, Germany) following the manufacturer’s protocol. The cDNA samples with Quantinova SYBR Green PCR kit (Qiagen, Germany) were prepared in a final volume of 20 μL and then subjected to RT-qPCR using LightCycler® 480 instrument (Roche Diagnostics). qPCR reaction was performed under the following conditions: 95°C for 2 minutes for pre-denaturing step, followed by 40 cycles of denaturation step at 95°C for 5 seconds, annealing at 60°C for 30 seconds, extension at 72°C for 30 seconds and finished with a cooling step at 4°C for 2 minutes. Primers for these three unigenes were purchased from Qiagen and each sample were performed in triplicates. The mRNA expression of all genes was quantified using 2^−ΔΔCT^ method with housekeeping genes *GAPDH* and *TBP*.

### Statistical analysis

Data analysis was performed using R/Bioconductor and Seaborn library in Python. Results of differentially expressed genes was visualised using EhancedVolcano package in R/Bioconductor. For gene expression studies, the fold change statistical significance was determined using One-way statistical analysis of variance (ANOVA). Statistical analysis details are described in the figure legends. All data are expressed as mean ± SD (Standard Deviation) (n = 3). To determine if datasets show a normal distribution, the Shapiro-Wilk test was used and P-values of less than 0.05 were considered as statistically significant.

## Acknowledgement

This study was supported by the Ministry Energy, Science, Technology, Environment and Climate Change (MESTECC) Malaysia Flagship Fund, reference number: FP0514B0021-2(DSTIN).

## Author contributions

A.V., U.A., S.C.C., D.M.C., S.L.C., S.A., and Y.K., designed the study. A.T performed the qPCR experiments. U.A. performed the wet and dry lab work as well as analysed the data. U.A wrote the paper. All authors reviewed the manuscript.

## Availability of data and materials

All data supporting the results in this article are included in the present and supplementary files except the raw and processed sequencing data that have been submitted to the NCBI Sequence Read Archive with accession number: PRJNA543209 (https://www.ncbi.nlm.nih.gov/bioproject/PRJNA543209) and Gene Expression Omnibus (GEO) with accession number GSE140902 (https://www.ncbi.nlm.nih.gov/geo/query/acc.cgi?acc=GSE140902).

## Ethics approval and consent to participate

Not application.

## Consent for publication

Not applicable.

