## Supplementary files for "Functional Transcriptome Analysis of Bladder Cancer Cell Lines Persistently Infected with Oncolytic Newcastle Disease Virus"

**Supplementary information**

### Mapping reads to a reference genome

While carrying out the quality control of the raw reads, adapter sequences were removed and ribosomal RNA (rRNA) sequences contamination were filtered to ensure that only the clean reads counts was used for mapping. The reference human DNA sequence file in FASTA format downloaded from Ensembl genome browser [1] was used to map each paired-end reads (in FASTQ format) to the GRCh37 (known as hg19) human reference genome using HISAT2, a faster and sensitive aligner that is suitable for splice site alignment[2]. As shown in Supplementary Table 1, the overall mapping rate for all the samples are above 95%, suggesting that majority of the paired-end reads are properly mapped to the human reference genome (Ensembl *GRCh37.p13*). Furthermore, the mapping rate was summarized by type to gain insight into which sample was properly mapped and which was not. The results yielded samples that were uniquely mapped, discordantly uniquely mapped, one mate mapped uniquely, multimapped, one mate multimapped and neither mate aligned (Supplementary figure 1).

Supplementary Table 1: Alignment rate for each of the RNA-seq read against the hg19 reference genome

| **S/No.** | **Samples ID** | **Sample Name** | **% Aligned** |
| --- | --- | --- | --- |
| 1 | ndv11_S16_L006 | EJ28Pi_R1 | 97.2% |
| 2 | ndv12_S17_L006 | EJ28Pi_R2 | 97.2% |
| 3 | ndv13_S18_L007 | EJ28Pi_R3 | 97.2% |
| 4 | ndv14_S19_L007 | EJ28_R1 | 96.7% |
| 5 | ndv15_S20_L007 | EJ28_R2 | 96.9% |
| 6 | ndv16_S21_L007 | EJ28_R3 | 97.1% |
| 7 | ndv17_S22_L007 | TCCSUPPi_R1 | 96.5% |
| 8 | ndv18_S23_L007 | TCCSUPPi_R2 | 96.3% |
| 9 | ndv19_S10_L003 | TCCSUPPi_R3 | 95.9% |
| 10 | ndv19_S43_L003 | TCCSUPPi_R3 | 95.7% |
| 11 | ndv20_S11_L003 | TCCSUP_R1 | 96.5% |
| 12 | ndv20_S44_L003 | TCCSUP_R1 | 96.4% |
| 13 | ndv21_S12_L003 | TCCSUP_R2 | 97.2% |
| 14 | ndv21_S37_L002 | TCCSUP_R2 | 97.5% |
| 15 | ndv22_S13_L003 | TCCSUP_R3 | 97.0% |
| 16 | ndv22_S45_L003 | TCCSUP_R3 | 97.0% |

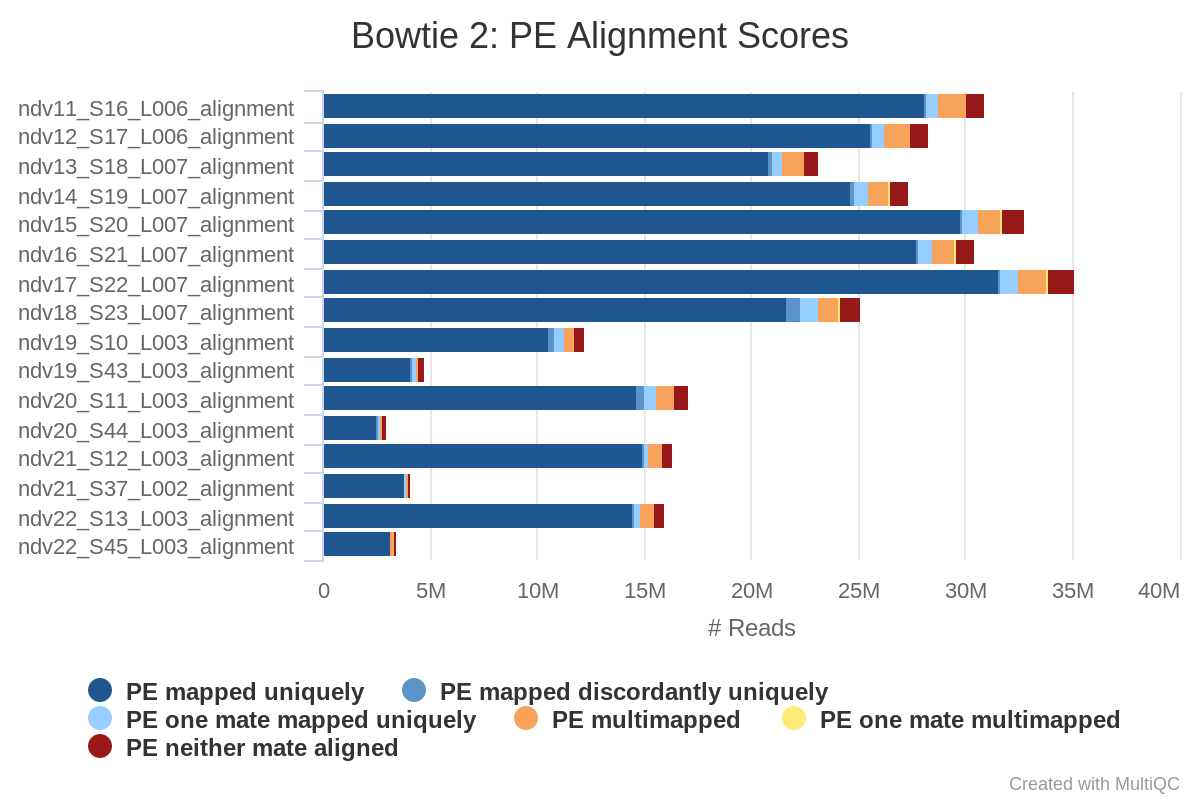

Supplementary Figure 1: HISAT2 read mapping by type

### Quantification of gene expression

Following the mapping of the RNA-Seq reads to the reference genome, transcripts abundance (measure of gene expression) was quantified using Salmon’s quasimapping strategy [3] that counts the number of reads that map to a given transcript. To ensure that only protein coding genes were captured for differential expression analysis (DEGs), a reference file that contains all the annotated human protein coding genes of known cDNA and ncRNA were downloaded from Ensembl’s website [1]. This file was then used to map each paired-end reads to these cDNA regions of interest and the transcript abundance estimation (features counts) was rapidly generated with all samples having a mean fragment length distribution peak of about 200 bp (supplementary figure 2), demonstrating that the transcripts expression level will be accurately estimated in all the samples. The mappings for each sample was subsequently used in DESeq2 package [4] via tximport for further analysis in R environment[5].

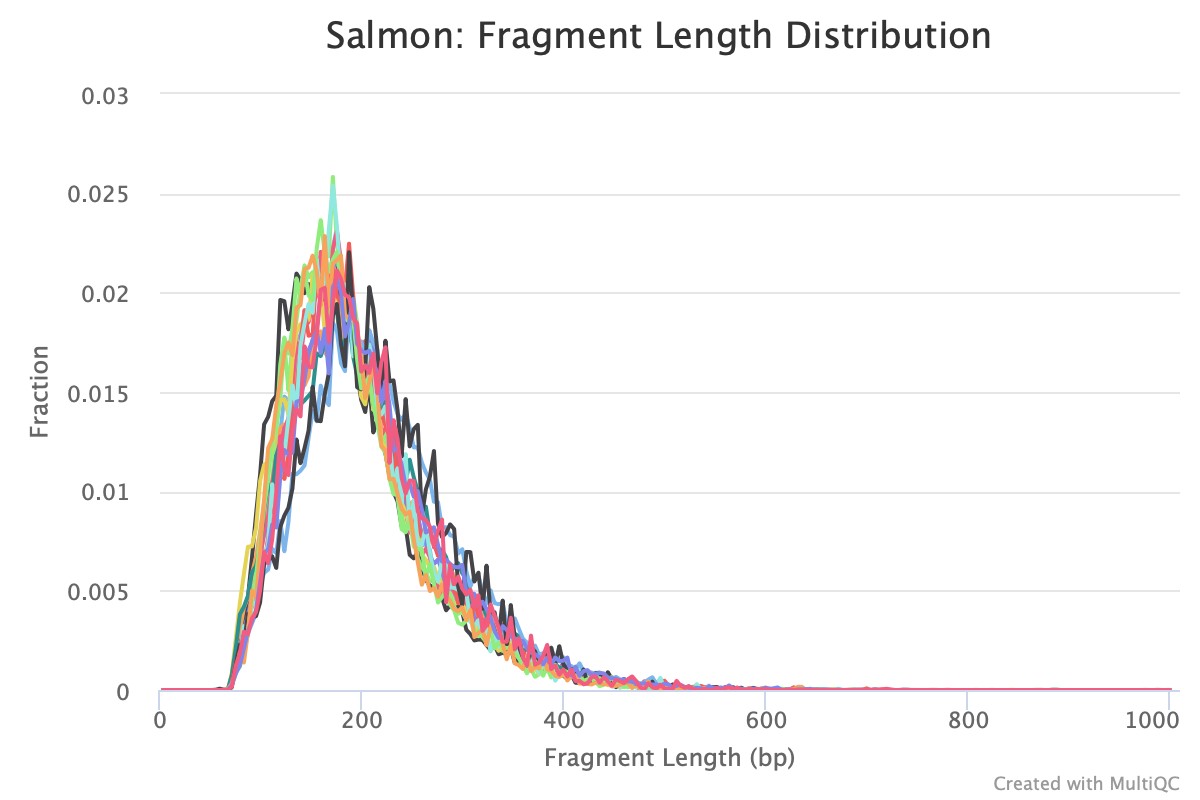

Supplementary Figure 2: Fragment length distribution of transcript from each read file based on quasimapping.

**Supplementary Figure 3: Principal component analysis (PCA) for all the RNA-Seq samples.** A) PCA plot the overall gene expression that separate TCCSUPPi cells from TCCSUP cells. B) PCA plot the overall gene expression that separate EJ28Pi cells from EJ28 cells.

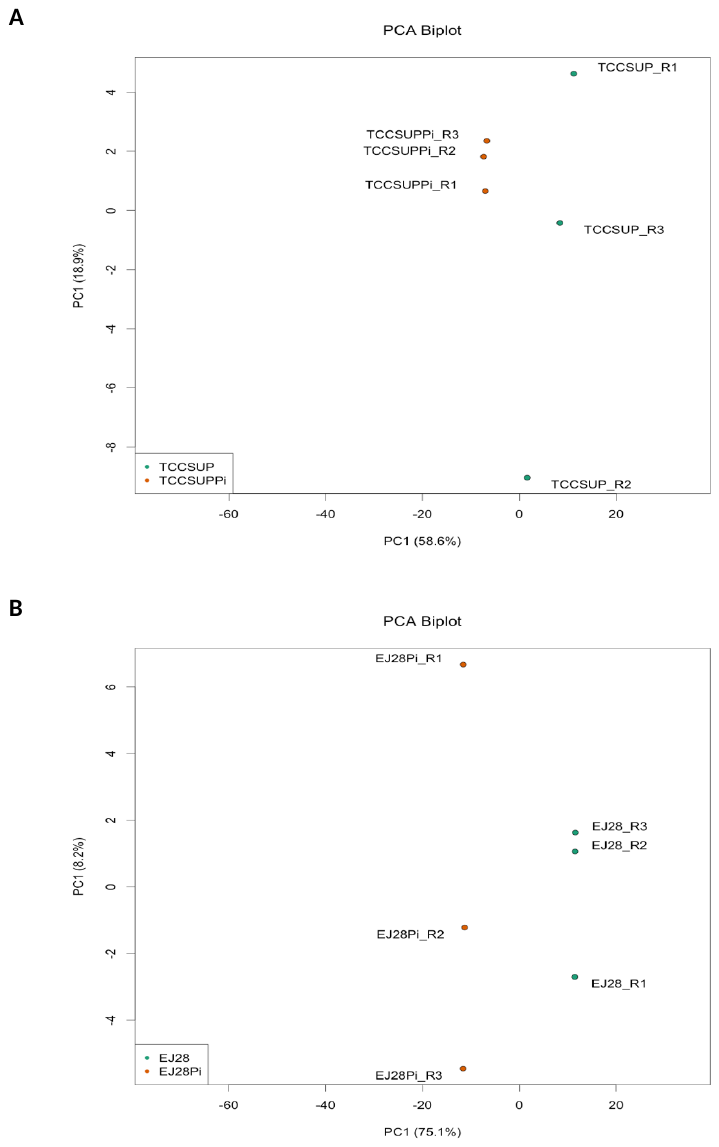

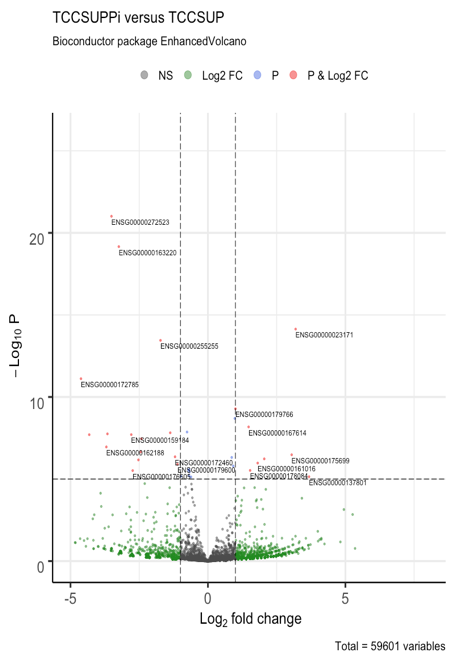

**A**

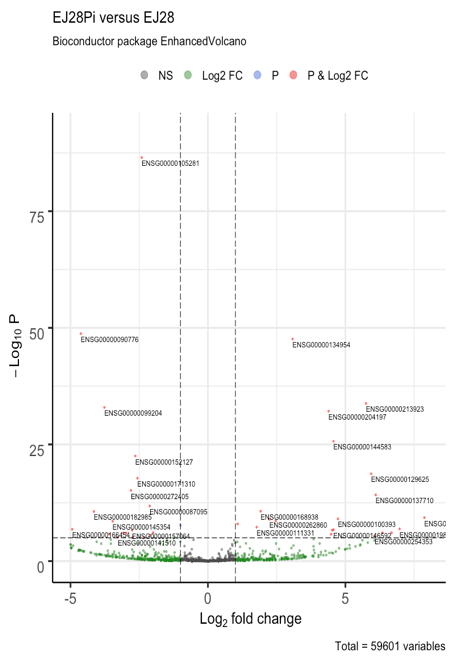

**B**

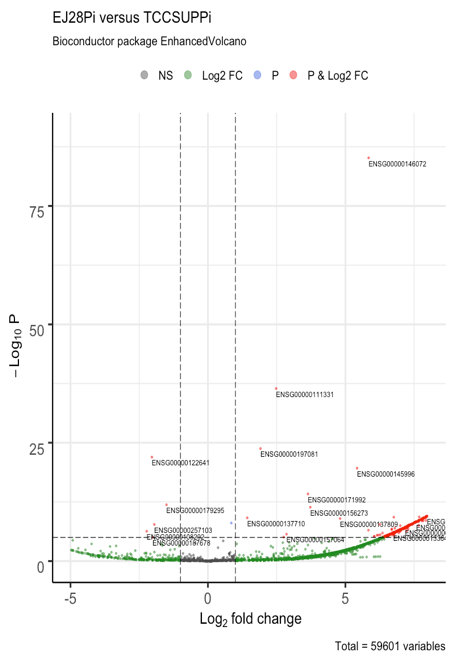

**C**

**Supplementary Fig. 4: Volcano plot of differentially expressed genes in persistently infected cancer cells and control cells.** A) Volcano plot for TCCSUPPi versus TCCSUP libraries. Profiles the relationship between the raw *p* values and log_2_ fold change in normalized expression (FPKM) between TCCSUPPi and TCCSUP. B) Volcano plot for EJ28Pi versus EJ28 libraries. Profiles the relationship between the raw *p* values and log_2_ fold change in normalized expression (FPKM) between EJ28Pi and EJ28. C) Volcano plot for EJ28Pi versus TCCSUPPi libraries. Profiles the relationship between the raw p values and log_2_ fold change in normalized expression (FPKM) between EJ28Pi and TCCSUPPi. Data is based on normalized FPKM (abundance) for each gene for each sample.

Supplementary Table 2: The enriched gene ontology (GO) categories of DEGs in EJ28Pi relative to TCCSUPPi

| **Category** | **GO ID** | **GO term** | **No. of DEGs** | **Log10**  **(*p* value)** | **DEG(s)** |
| --- | --- | --- | --- | --- | --- |
| Biological process | GO:0044843 | Cell cycle G1/S phase transition | 21 | -3.5188 | *ANXA1,BACH1,CDC6,CDKN2B,EP300,GFI1,*  *SFN,GSPT1,INHBA,MNAT1,PLRG1,RDX,SOX4,*  *CUL3,CUL1,BRD4,BRD7,RPS27L,CNOT6,SPDYA,STXBP4* |
|  | GO:0000082 | G1/S transition of mitotic cell cycle | 20 | -3.4748 | *ANXA1,BACH1,CDC6,CDKN2B,EP300,GFI1,SFN,*  *GSPT1,INHBA,MNAT1,PLRG1,RDX,SOX4,CUL3,CUL1,*  *BRD4,BRD7,RPS27L,CNOT6,SPDYA* |
|  | GO:0007169 | Transmembrane receptor protein tyrosine kinase signalling pathway | 38 | -3.1876 | *ACTB,AP2A1,AXL,CEACAM1,CDC42,AP3S1,COL1A1,*  *DCN,EFNB1,EPHB2,EPHB3,IFI6,GRB2,HGF,HNRNPH1,*  *IGF2,IGF2R,IGFBP5,IRS1,PIGR,PTPN11,PTPRA,*  *PTPRR,ROS1,SREBF1,THBS1,TYRO3,FGF17,GPRC5A*  *,MAPKAPK2,CEP57,CD3EAP,SLC30A10,TMEM204*  *,SPRY4,FAM83A,STXBP4,TRIM72* |
|  | GO:0038171 | Cannabinoid signalling pathway | 3 | -3.0599 | *DAGLA,CNR2,MGLL* |
|  | GO:0044770 | Cell cycle phase transition | 33 | -2.7460 | *ANXA1,BACH1,CDC6,CDKN2B,CSNK1E,EP300,*  *GFI1,SFN,GSPT1,INHBA,MNAT1,OVOL1,PLRG1,*  *PPP1CB,RDX,SOX4,ZNF207,CUL3,CUL1,CEP57,*  *TACC3,CEP131,BRD4,BRD7,RPS27L,MRNIP,HAUS6,*  *CNOT6,CCNJL,DBF4B,NEK10,SPDYA,STXBP4* |
|  | GO:0010564 | Regulation of cell cycle process | 39 | -2.7339 | *ANXA1,BCL2L1,CDC6,CDC42,CDKN2B,CSNK1E,DAZL,*  *EP300,GNAI1,SFN,IGF2,LRP5,OVOL1,PDE3A,PLRG1,*  *RAD51,RDX,SOX4,ZNF207,SMC1A,CUL3,CUL1,CEP57,*  *TACC3,GIPC1,CBX3,CEP131,BRD4,BRD7,RPS27L,MRNIP,*  *HAUS6,KLHL9,CNOT6,ANKRD53,DBF4B,SLF1,NEK10,STXBP4* |
|  | GO:0031623 | Receptor internalization | 10 | -2.6822 | *ACHE,CEACAM1,CD36,GRB2,LRPAP1,SELE,SNX1,*  *PICALM,RAMP1,ANKRD13D* |
|  | GO:0051301 | Cell division | 31 | -2.6090 | *BCL2L1,CDC6,CDC42,CFL1,GNAI1,SFN,IGF2,KIFC1,*  *LLGL2,MAP4,MYH9,PPP1CB,ZNF207,SMC1A,CUL3,*  *RUVBL1,NCAPD2,TACC3,GIPC1,TXNL4A,TRIOBP,*  *CKAP2,ALKBH4,HAUS6,INTS13,KLHL9,PRDM15,*  *ANKRD53,PARD6G,CENPX,REEP3,TACC2* |
|  | GO:0007346 | Regulation of mitotic cell cycle | 33 | -2.4539 | *ANXA1,BCL2L1,CDC6,CDKN2B,CSNK1E,EP300,*  *GNAI1,SFN,IGF2,LRP5,OVOL1,PLRG1,PTPN11,*  *RDX,SOX4,ZNF207,SMC1A,CUL3,CUL1,CEP57,*  *TACC3,CEP131,BRD4,BRD7,RPS27L,MRNIP,HAUS6,*  *INTS13,CNOT6,ANKRD53,DBF4B,MUS81,SLF1* |
|  | GO:0045787 | Positive regulation of cell cycle | 22 | -2.4221 | *ANXA1,CDC6,CDC42,DAZL,EP300,SFN,IGF2,LRP5,*  *MNAT1,OVOL1,PLRG1,PTPN11,RDX,SOX4,CUL3,*  *GIPC1,BRD4,CNOT6,DBF4B,SLF1,SPDYA,STXBP4* |
|  | GO:0044772 | Mitotic cell cycle phase transition | 30 | -2.3911 | *ANXA1,BACH1,CDC6,CDKN2B,CSNK1E,EP300,*  *GFI1,SFN,GSPT1,INHBA,MNAT1,PLRG1,PPP1CB,*  *RDX,SOX4,ZNF207,CUL3,CUL1,CEP57,TACC3,CEP131,*  *BRD4,BRD7,RPS27L,MRNIP,HAUS6,CNOT6,CCNJL,*  *DBF4B,SPDYA* |
|  | GO:0090068 | Positive regulation of cell cycle process | 18 | -2.3825 | *ANXA1,CDC6,CDC42,DAZL,EP300,SFN,*  *IGF2,LRP5,PLRG1,RDX,SOX4,CUL3,GIPC1,*  *BRD4,CNOT6,DBF4B,SLF1,STXBP4* |
|  | GO:0006400 | tRNA modification | 8 | -2.3564 | *HSD17B10,METTL1,TARBP1,THUMPD3,*  *CDKAL1,TRMT10C,OSGEP,CTU2* |
|  | GO:0036465 | Synaptic vesicle recycling | 7 | -2.3288 | *ACTB,SYT1,PICALM,SYT7,SNAP91,GRIPAP1,BTBD9* |
|  | GO:0034498 | Early endosome to Golgi transport | 3 | -2.3105 | *SNX1,SNX2,STX5* |
|  | GO:0051592 | Response to calcium ion | 11 | -2.2322 | *FGB,GUCA1A,HSPA5,MNAT1,SYT1,*  *THBS1,SYT7,TRPV6,SUCNR1,SYT13,SYT15* |
|  | GO:0030488 | tRNA methylation | 5 | -2.2186 | *HSD17B10,METTL1,TARBP1,THUMPD3,TRMT10C* |
|  | GO:0008033 | tRNA processing | 10 | -2.2175 | *HSD17B10,METTL1,TARBP1,ADAT1,THUMPD3,*  *CDKAL1,RPP25,TRMT10C,OSGEP,CTU2* |
|  | GO:0070525 | tRNA threonylcarbamoyladenosine metabolic process | 3 | -2.2063 | *HSD17B10,TRMT10C,OSGEP* |
|  | GO:0030677 | Ribonuclease P complex | 3 | -2.1113 | *HSD17B10,RPP25,TRMT10C* |
| Cellular component | GO:0030133 | Transport vesicle | 25 | -3.3771 | *AP2A1,BCL2L1,CEACAM1,AP3S1,IGF2R,LAMP1,NTS,PAM,SREBF1,*  *STX5,SYPL1,SYT1,SLC30A3,PICALM,*  *SYT7,SEC24D,SNAP91,GIPC1,COPG2*  *,SYT13,SPX,SYT15,ZNRF1,RAB3C,SYCN,ANXA1*  *,CD14,MS4A3,CD36,CLCA1,B4GALT1,GRB2,PIGR,*  *GPRC5A,SNAP29,NMNAT2,*  *KIF1B,NECAP1,TBC1D24,CLEC4D,RAB43,TRIM72* |
|  | GO:0030117 | Membrane coat | 10 | -3.1967 | *AP2A1,AP3S1,IGF2R,SCN10A,PICALM,*  *SEC24D,NECAP1,COPG2,AP5M1,SCLT1* |
|  | GO:0048475 | Coated membrane | 10 | -3.1967 | *AP2A1,AP3S1,IGF2R,SCN10A,PICALM,*  *SEC24D,NECAP1,COPG2,AP5M1,SCLT1* |
|  | GO:0070161 | Anchoring junction | 29 | -2.4176 | *ACTB,ANXA1,CEACAM1,CDC42,CDH2,CDH5,*  *CDH8,CFL1,CNN2,DSC3,DSG2,B4GALT1,HSPA5,IGF2R,*  *ITGB4,MYH9,PPP1CB,RDX,RPL8,RPL38,LIMD1,TRIOBP,*  *ITGA11,LIN7C,CORO1B,TMEM204,ARMC5,SPRY4,DSG4* |
|  | GO:0030658 | Transport vesicle membrane | 15 | -2.7146 | *AP2A1,BCL2L1,CEACAM1,LAMP1,PAM,SREBF1,*  *STX5,SYPL1,SYT1,SLC30A3,SYT7,SEC24D,ZNRF1,*  *RAB3C,SYCN* |
|  | GO:0012506 | Vesicle membrane | 37 | -2.2098 | *AP2A1,ANXA1,BCL2L1,CEACAM1,CD14,MS4A3,CD36,*  *AP3S1,CLCA1,B4GALT1,GRB2,IGF2R,LAMP1,PAM,PIGR,*  *SREBF1,STX5,SYPL1,SYT1,SLC30A3,PICALM,GPRC5A,SYT7,*  *SNAP29,SEC24D,GIPC1,NMNAT2,KIF1B,NECAP1,COPG2,*  *TBC1D24,ZNRF1,RAB3C,CLEC4D,RAB43,SYCN,TRIM72* |
|  | GO:1902555 | Endoribonuclease complex | 5 | -2.6628 | *HSD17B10,PRKRA,AGO1,RPP25,TRMT10C* |
|  | GO:1905348 | Endonuclease complex | 5 | -2.5997 | *HSD17B10,PRKRA,AGO1,RPP25,TRMT10C* |
|  | GO:0030677 | Ribonuclease P complex | 3 | -2.1113 | *HSD17B10,RPP25,TRMT10C* |
|  | GO:0030118 | Clathrin coat | 6 | -2.6086 | *AP2A1,IGF2R,SCN10A,PICALM,NECAP1,SCLT1* |
|  | GO:0005667 | Transcription factor complex | 22 | -2.7876 | *CTBP1,EP300,ETS1,SMAD2,MNAT1,NFATC1,PBX2,POU2F1,*  *SNAPC1,SOX4,SRY,TCF12,THRA,FOXH1,LIMD1,HDAC9,*  *HOXB13,NFAT5,CD3EAP,HES6,ZFPM1,GTF2H2C* |
|  | GO:0044798 | Nuclear transcription factor complex | 13 | -2.1223 | *SMAD2,MNAT1,NFATC1,POU2F1,SNAPC1,*  *SOX4,SRY,TCF12,THRA,FOXH1,NFAT5,*  *CD3EAP,GTF2H2C* |
|  | GO:0030863 | Cortical cytoskeleton | 10 | -2.6536 | *ACTB,CDH2,CFL1,EEF1A1,LLGL2,MYH9,RDX,SELE,*  *SPTAN1,MYADM,CTBP1,GNAI1* |
|  | GO:0030864 | Cortical actin cytoskeleton | 8 | -2.3875 | *CDH2,CFL1,EEF1A1,LLGL2,MYH9,RDX,SPTAN1,MYADM* |
|  | GO:0044448 | Cell cortex part | 12 | -2.0469 | *ACTB,CDH2,CFL1,CTBP1,EEF1A1,GNAI1,LLGL2,*  *MYH9,RDX,SELE,SPTAN1,MYADM* |
| Molecular function | GO:0005229 | Intracellular calcium activated chloride channel activity | 6 | -5.1653 | *CLCA1,CLCA4,ANO2,TTYH1,ANO8,ANO9,GABRE* |
|  | GO:0001786 | Phosphatidylserine binding | 9 | -4.137 | *AXL,SYT1,THBS1,SYT7,GSDMC,GRAMD1B,SYT13,SYT15,TRIM72* |
|  | GO:0050839 | Cell adhesion molecule binding | 31 | -3.9419 | *ANXA1,CDH2,CDH5,CDH8,CNN2,DSG2,FGB,SFN,*  *HSPA5,IGF2,ITGB4,MYH9,PTPN11,RDX,RPL29,SNX1,*  *SNX2,SPTAN1,STX5,THBS1,PICALM,TAGLN2,RUVBL1,*  *PROM1,H1FX,GPRC5A,CD226,GIPC1,ADAMTS8,SERBP1,CORO1B* |
|  | GO:0000149 | SNARE binding | 12 | -3.9156 | *STX5,SYPL1,SYT1,PICALM,SYT7,SNAP29,SEC24D,*  *SNAP91,GABARAPL2,SYT13,SYT15,STXBP4* |
|  | GO:0030276 | clathrin binding | 8 | -3.5324 | *AP2A1,SYT1,PICALM,SYT7,SNAP91,SYT13,SYT15,SCLT1* |
|  | GO:0005158 | Insulin receptor binding | 5 | -3.3753 | *IGF2,IRS1,PTPN11,SNX1,SNX2,STX5* |
|  | GO:0016423 | tRNA (guanine) methyltransferase activity | 4 | -3.3273 | *METTL1,TARBP1,THUMPD3,TRMT10C* |
|  | GO:0019900 | Kinase binding | 38 | -3.1098 | *ACTB,AP2A1,BCL2L1,CEACAM1,CDC6,CDC42,CDH2,*  *CDKN2B,EEF1A1,SFN,GRB2,IRS1,MYCN,NFATC1,PAM*  *,PARN,PDE8A,PPP1CB,PTPN11,PTPRR,SREBF1,TCL1A,*  *MAPKAPK2,HDAC9,SNAP91,CD226,KIF1B,PPP1R15A,*  *NBEA,CDK12,BANK1,CCNJL,SIKE1,DBF4B,PARD6G,FAM83A,*  *SPDYA,ALKAL2* |
|  | GO:0019901 | Protein kinase binding | 34 | -2.8474 | *ACTB,AP2A1,BCL2L1,CEACAM1,CDC42,CDH2,CDKN2B*  *,EEF1A1,SFN,GRB2,IRS1,NFATC1,PAM,PARN,PPP1CB,*  *PTPN11,PTPRR,SREBF1,TCL1A,MAPKAPK2,HDAC9,SNAP91,*  *CD226,PPP1R15A,NBEA,CDK12,BANK1,CCNJL,SIKE1,DBF4B,*  *PARD6G,FAM83A,SPDYA,ALKAL2* |
|  | GO:0000978 | RNA polymerase II proximal promoter sequence-specific DNA binding | 29 | -2.7471 | *ACTB,KLF6,EP300,ETS1,SMAD2,MYCN,NFATC1,NFIC,*  *OTX1,OVOL1,POU2F1,SNAI2,SOX4,SP3,SREBF1,STAT2,*  *TCF12,NR2F1,BHLHE40,TBX19,ARID3B,NFAT5,NKX2-8,AGO1,KLF15,PRDM15,ZFHX2,AEBP2,ZFPM1,BACH1,*  *PBX2,SATB1,THRA,IRF2BP1,PURG,HES6,IRF2BPL,*  *FOXK1,FOXH1,ZFAT* |
|  | GO:0072341 | Modified amino acid binding | 9 | -2.7381 | *AXL,SYT1,THBS1,SYT7,GSDMC,GRAMD1B,SYT13,SYT15,TRIM72* |
|  | GO:0008134 | Transcription factor binding | 33 | -2.6174 | *ACTB,CTBP1,EP300,ETS1,HDGF,IFI27,IGHMBP2,*  *SMAD2,NFATC1,PBX2,PTMA,TRAPPC2,SRY,TCF12,*  *THRA,VHL,BHLHE40,RUVBL1,FOXH1,VGLL4,HDAC9*  *,MED24,TACC2,NFAT5,BRD4,ANKRD1,BRD7,*  *NLK,CDK12,WIPI1,HES6,HMGB4,ZFPM1* |
|  | GO:0001228 | DNA-binding transcription activator activity, RNA polymerase II-specific | 24 | -2.5831 | BACH1,KLF6,EP300,ETS1,SMAD2,MYCN,NFATC1,NFIC,OTX1  ,OVOL1,PBX2,SOX4,SREBF1,STAT2,TCF12,FOXH1,TBX19,  ARID3B,NFAT5,IRF2BP1,NKX2-8,KLF15,ZFAT,IRF2BPL |
|  | GO:0000987 | Proximal promoter sequence-specific DNA binding | 29 | -2.5608 | *ACTB,KLF6,EP300,ETS1,SMAD2,MYCN,NFATC1,NFIC,OTX1,*  *OVOL1,POU2F1,SNAI2,SOX4,SP3,SREBF1,STAT2,TCF12,*  *NR2F1,BHLHE40,TBX19,ARID3B,NFAT5,NKX2-8,AGO1,KLF15,PRDM15,ZFHX2,AEBP2,ZFPM1* |
|  | GO:0001012 | RNA polymerase II regulatory region DNA binding | 38 | -2.5521 | *ACTB,BACH1,KLF6,EP300,ETS1,SMAD2,MYCN,NFATC1,*  *NFIC,OTX1,OVOL1,PBX2,POU2F1,SATB1,SNAI2,SOX4,*  *SP3,SREBF1,STAT2,TCF12,NR2F1,THRA,BHLHE40,TBX19,*  *ARID3B,NFAT5,IRF2BP1,NKX28,AGO1,KLF15,PURG,*  *HES6,PRDM15,IRF2BPL,ZFHX2,AEBP2,ZFPM1,FOXK1* |
|  | GO:0050750 | Low-density lipoprotein particle receptor binding | 4 | -2.4878 | *AP2A1,LRPAP1,SYT1,PICALM* |
|  | GO:0008289 | Lipid binding | 36 | -2.4082 | *ADH5,ALOX15B,ANXA1,AXL,BTK,CD14,CD36,CRABP2,F2,*  *GPR12,IGF2R,RAG2,S100A9,SNX1,SNX2,SYT1,NR2F2,*  *THBS1,PICALM,SOAT2,PROM1,SYT7,SNAP91,PEMT,*  *PLEK2,EPDR1,WIPI1,GSDMC,GRAMD1B,SYT13,SYT15,*  *STARD3NL,BPIFA3,BPIFA2,OXER1,TRIM72* |
|  | GO:0070325 | Lipoprotein particle receptor binding | 4 | -2.1419 | *AP2A1,LRPAP1,SYT1,PICALM* |
|  | GO:0005254 | Chloride channel activity | 7 | -2.1309 | *CLCA1,GABRE,CLCA4,ANO2,TTYH1,ANO8,ANO9* |
|  | GO:0000049 | tRNA binding | 6 | -2.1047 | *EEF1A1,HSD17B10,IGHMBP2,METTL1,TRMT10C,CTU2* |

Supplementary Table 3: The enriched gene ontology (GO) categories of DEGs in TCCSUPPi relative to control (TCCSUP)

| **Category** | **GO ID** | **GO term** | **No. of DEGs** | **Log10**  **(*p* value)** | **DEG(s)** |
| --- | --- | --- | --- | --- | --- |
| Biological process | GO:0006986 | Response to unfolded protein | 5 | -4.3345 | *CALR,HSPA1B,HSPA4,THBS1,HSPB7,*  *HOXB13,NUDCD2,NR2F1,GPHB5* |
|  | GO:0035966 | Response to topologically incorrect protein | 5 | -4.0882 | *CALR,HSPA1B,HSPA4,THBS1,HSPB7* |
|  | GO:0033574 | Response to testosterone | 3 | -3.9832 | *CALR,THBS1,HOXB13* |
|  | GO:0060326 | Cell chemotaxis | 4 | -2.2903 | *CALR,S100A9,THBS1,HMGB4* |
|  | GO:0032103 | Positive regulation of response to external stimulus | 4 | -2.1801 | *CALR,F2,S100A9,THBS1* |
|  | GO:0030522 | Intracellular receptor signalling pathway | 4 | -2.4215 | *CALR,HSPA1B,NR2F1,GPHB5* |
|  | GO:0006626 | Protein targeting to mitochondrion | 3 | -2.8364 | *HSPA4,TOMM7,UBL4B,RPL8,ZDHHC7* |
|  | GO:0006605 | Protein targeting | 5 | -2.5253 | *HSPA4,RPL8,TOMM7,ZDHHC7,UBL4B* |
|  | GO:0072655 | Establishment of protein localization to mitochondrion | 3 | -2.4232 | *HSPA4,TOMM7,UBL4B* |
|  | GO:0070585 | Protein localization to mitochondrion | 3 | -2.3882 | *HSPA4,TOMM7,UBL4B* |
|  | GO:0034332 | Adherens junction organization | 3 | -2.3796 | *CDH2,CDH5,THBS1* |
|  | GO:0031331 | Positive regulation of cellular catabolic process | 4 | -2.0367 | *HSPA1B,TOMM7,TMEM259,HMGB4* |
| Cellular component | GO:0062023 | Collagen-containing extracellular matrix | 7 | -4.4250 | *BGN,CALR,CDH2,F2,S100A9,THBS1,PODNL1,*  *CDH5,ACER3,TTYH1,LRPAP1,HMGB4* |
|  | GO:0031012 | Extracellular matrix | 7 | -3.7271 | *BGN,CALR,CDH2,F2,S100A9,THBS1,PODNL1* |
|  | GO:0005788 | Endoplasmic reticulum lumen | 5 | -3.1653 | *CALR,CDH2,F2,LRPAP1,THBS1* |
|  | GO:0005796 | Golgi lumen | 3 | -2.7865 | *BGN,F2,LRPAP1* |
|  | GO:0019897 | Extrinsic component of plasma membrane | 3 | -2.1987 | *CDH2,CDH5,GNG3* |
|  | GO:0005912 | Adherens junction | 5 | -2.0960 | *CALR,CDH2,CDH5,HSPA1B,RPL8* |
| Molecular function | GO:0005509 | Calcium ion binding | 8 | -3.73258 | *CALR,CDH2,CDH5,F2,S100A9,THBS1,ACER3,TTYH1* |
|  | GO:0001227 | DNA-binding transcription repressor activity, RNA polymerase II-specific | 5 | -3.68349 | *BACH1,NR2F1,HOXB13,HES6,AEBP2* |
|  | GO:0005539 | Glycosaminoglycan binding | 4 | -2.7221 | *BGN,F2,LRPAP1,THBS1* |
|  | GO:0008201 | Heparin binding | 3 | -2.1700 | *F2,LRPAP1,THBS1* |
|  | GO:0051082 | Unfolded protein binding | 3 | -2.47778 | *CALR,HSPA1B,NUDCD2* |

Supplementary Table 4: The enriched gene ontology (GO) categories of DEGs in EJ28Pi relative to control (EJ28)

| **Category** | **GO ID** | **GO term** | **No. of DEGs** | **Log10**  **(*p* value)** | **DEG(s)** |
| --- | --- | --- | --- | --- | --- |
| Biological process | GO:0008285 | Negative regulation of cell proliferation | 17 | -6.0428 | *ETS1,ETV3,SFN,INHBA,MNT,MSX1,EIF2AK2,*  *PROX1,ZEB1,TP53,WNT9A,WT1,PTGES,P3H2,CAMK2N1,*  *NDRG4,KCTD11,FGF9,IGF2,SYT1,EI24,BASP1,FASLG,UNC5B* |
|  | GO:0060322 | Head development | 17 | -6.0118 | *CSNK1E,EP300,ETS1,FGF9,ARHGAP35,INHBA,MSX1,*  *PITX1,PPP3CA,PROX1,PTPN11,RORA,SYT1,THRA,*  *TP53,BASP1,NDRG4,ZEB1,ANKRD1,KCTD11,RAC1* |
|  | GO:0007420 | Brain development | 16 | -5.6634 | *CSNK1E,ETS1,FGF9,ARHGAP35,INHBA,MSX1*  *,PITX1,PPP3CA,PROX1,PTPN11,RORA,SYT1,THRA,TP53,*  *BASP1,NDRG4* |
|  | GO:0000075 | Cell cycle checkpoint | 9 | -5.5456 | *CCND1,FOXN3,EP300,SFN,PROX1,*  *PTPN11,TP53,WEE1,WNT9A* |
|  | GO:0010906 | Regulation of glucose metabolic process | 7 | -5.4501 | *EP300,IGF2,IRS1,PPP1CB,RORA,TP53,FOXK1* |
|  | GO:0048598 | Embryonic morphogenesis | 14 | -5.4379 | *FGF9,ARHGAP35,IGF2,INHBA,AFF3,MSX1,PITX1,*  *PROX1,ZEB1,TP53,WNT9A,CHST11,LMBR1,NDRG4,*  *PDGFA,THRA,WT1,FOXN3,EP300,PTPN11,FASLG* |
|  | GO:0030326 | Embryonic limb morphogenesis | 7 | -5.2594 | *FGF9,AFF3,MSX1,PITX1,WNT9A,CHST11,LMBR1* |
|  | GO:0048732 | Gland development | 12 | -5.3074 | *CCND1,ETS1,ARHGAP35,IGF2,MSX1,PDGFA,PITX1,*  *PROX1,STAT6,THRA,WT1,LBH* |
|  | GO:0048511 | Rhythmic process | 10 | -5.2518 | *CSNK1E,EP300,ETS1,INHBA,PPP1CB,PROX1,*  *RORA,SP1,TP53,BHLHE40,CCND1,NDRG4* |
|  | GO:0010675 | Regulation of cellular carbohydrate metabolic process | 7 | -4.8958 | *EP300,IGF2,IRS1,PPP1CB,RORA,TP53,FOXK1* |
|  | GO:0035107 | Appendage morphogenesis | 7 | -4.7789 | *FGF9,AFF3,MSX1,PITX1,WNT9A,CHST11,LMBR1* |
|  | GO:0007517 | Muscle organ development | 11 | -4.8013 | *DMD,EP300,FGF9,MSX1,PITX1,PPP3CA,PROX1,WT1,*  *BASP1,ANKRD1,FOXK1,IGF2,RORA,ZEB1,PDGFA,*  *THRA,PTPN11,SYT1,TP53,CHST11,NDRG4,*  *SFN,CCND1,INHBA,EFNB1* |
|  | GO:0044772 | Mitotic cell cycle phase transition | 13 | -4.6880 | *CCND1,FOXN3,CSNK1E,EP300,SFN,INHBA,*  *PPP1CB,PPP3CA,RDX,TP53,WEE1,CEP135,CTDSPL* |
|  | GO:0061448 | Connective tissue development | 9 | -4.7272 | *FGF9,MSX1,PDGFA,PITX1,ZEB1,THRA,WNT9A,*  *WT1,CHST11* |
|  | GO:0061061 | Muscle structure development | 14 | -4.7006 | *DMD,EP300,FGF9,IGF2,MSX1,PITX1,PPP3CA,*  *PROX1,RORA,ZEB1,WT1,BASP1,ANKRD1,FOXK1* |
|  | GO:0000082 | G1/S transition of mitotic cell cycle | 9 | -4.6528 | *CCND1,EP300,SFN,INHBA,PPP3CA,*  *RDX,TP53,WEE1,CTDSPL* |
|  | GO:0030031 | Cell projection assembly | 13 | -4.6047 | *CSNK1E,EMP2,ARHGAP35,ABLIM1,PDGFA,RAC1,*  *RDX,CEP135,ATMIN,CD2AP,RABGAP1,TBC1D16,SCLT1* |
|  | GO:0001501 | Skeletal system development | 12 | -4.5817 | *FOXN3,EP300,FGF9,IGF2,MSX1,PITX1,PTPN11,*  *ZEB1,THRA,TP53,WNT9A,CHST11* |
|  | GO:0046888 | Negative regulation of hormone secretion | 5 | -4.5585 | *INHBA,IRS1,PPP3CA,PTPN11,VSNL1* |
|  | GO:0007093 | Mitotic cell cycle checkpoint | 7 | -4.4749 | *CCND1,FOXN3,EP300,SFN,TP53,WEE1,WNT9A* |
| Cellular component | GO:0098589 | Membrane region | 9 | -4.0883 | *FASLG,DMD,EFNB1,EMP2,IRS1,OLR1,STAT6,XPO1,UNC5B* |
|  | GO:0045121 | Membrane raft | 8 | -3.4626 | *FASLG,DMD,EFNB1,EMP2,IRS1,OLR1,STAT6,UNC5B* |
|  | GO:0098857 | Membrane microdomain | 8 | -3.4535 | *FASLG,DMD,EFNB1,EMP2,IRS1,OLR1,STAT6,UNC5B* |
|  | GO:0005667 | Transcription factor complex | 7 | -2.4299 | *EP300,ETS1,PITX1,ZEB1,THRA,TP53,GTF2H2C* |
|  | GO:0005901 | Caveola | 3 | -2.0099 | *FASLG,EMP2,IRS1* |
| Molecular function | GO:0019904 | Protein domain specific binding | 19 | -8.0071 | *SFN,IRS1,PROX1,PTPN11,RDX,SP1,THRA,TP53,*  *WT1,XPO1,BHLHE40,VAPB,HOMER2,BASP1,*  *GIPC1,CD2AP,CADM1,NLK,PLEKHA2* |
|  | GO:0000977 | RNA polymerase II regulatory region sequence-specific DNA binding | 16 | -5.3173 | *EP300,ETS1,ETV3,ARHGAP35,MNT,MSX1,PITX1,*  *PROX1,RORA,SP1,STAT6,ZEB1,THRA,TP53,BHLHE40,*  *FOXK1,CCND1,ANKRD1,NLK,CDK12,WT1,WNT9A,GTF2H2C* |
|  | GO:0001012 | RNA polymerase II regulatory region DNA binding | 16 | -5.2681 | *EP300,ETS1,ETV3,ARHGAP35,MNT,MSX1,*  *PITX1,PROX1,RORA,SP1,STAT6,ZEB1,*  *THRA,TP53,BHLHE40,FOXK1* |
|  | GO:0008134 | Transcription factor binding | 14 | -4.941 | *CCND1,EP300,ETS1,PITX1,PROX1,RORA,SP1,*  *ZEB1,THRA,TP53,BHLHE40,ANKRD1,NLK,CDK12* |
|  | GO:0001227 | DNA-binding transcription repressor activity, RNA polymerase II-specific | 8 | -4.3169 | *ETV3,ARHGAP35,MNT,MSX1,PROX1,ZEB1,BHLHE40,FOXK1* |
|  | GO:0008022 | Protein C-terminus binding | 7 | -4.1451 | *FOXN3,EP300,SP1,SYT1,CEP135,CD2AP,SCLT1* |
|  | GO:0001085 | RNA polymerase II transcription factor binding | 6 | -3.9819 | *EP300,PITX1,SP1,TP53,BHLHE40,ANKRD1* |
|  | GO:0005242 | Inward rectifier potassium channel activity | 3 | -3.7797 | *KCNJ5,KCNJ10,KCNJ12* |
|  | GO:0005158 | Insulin receptor binding | 3 | -3.6525 | *IGF2,IRS1,PTPN11* |
|  | GO:0003714 | Transcription corepressor activity | 7 | -3.4966 | *CCND1,ARHGAP35,MNT,ZEB1,BHLHE40,BASP1,ANKRD1* |
|  | GO:0042826 | Histone deacetylase binding | 5 | -3.4540 | *CCND1,RAC1,SP1,TP53,ANKRD1* |
|  | GO:0002039 | P53 binding | 4 | -3.3383 | *EP300,MSX1,TP53,ANKRD1* |
|  | GO:0035035 | Histone acetyltransferase binding | 3 | -3.2893 | *ETS1,SP1,TP53* |
|  | GO:0001228 | DNA-binding transcription activator activity, RNA polymerase II-specific | 9 | -3.2755 | *EP300,ETS1,MSX1,PITX1,RORA,SP1,STAT6,TP53,WT1* |
|  | GO:0000978 | RNA polymerase II proximal promoter sequence-specific DNA binding | 10 | -3.1684 | *EP300,ETS1,PITX1,PROX1,RORA,SP1,STAT6*  *,ZEB1,TP53,BHLHE40* |
|  | GO:0000987 | Proximal promoter sequence-specific DNA binding | 10 | -3.0707 | *EP300,ETS1,PITX1,PROX1,RORA,SP1,STAT6,*  *ZEB1,TP53,BHLHE40* |
|  | GO:0008083 | Growth factor activity | 5 | -2.7041 | *FGF9,FGF14,IGF2,INHBA,PDGFA* |
|  | GO:0003712 | Transcription coregulator activity | 9 | -2.4530 | *CCND1,EP300,ARHGAP35,MNT,ZEB1,THRA,*  *BHLHE40,BASP1,ANKRD1* |
|  | GO:0030971 | Receptor tyrosine kinase binding | 3 | -2.2013 | *IRS1,PTPN11,TP53* |
|  | GO:0051219 | Phosphoprotein binding | 3 | -2.0098 | *SFN,IRS1,PTPN11* |

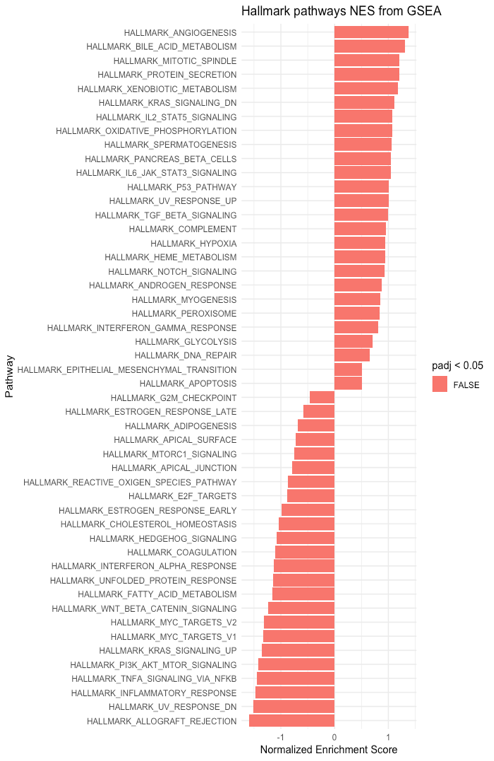

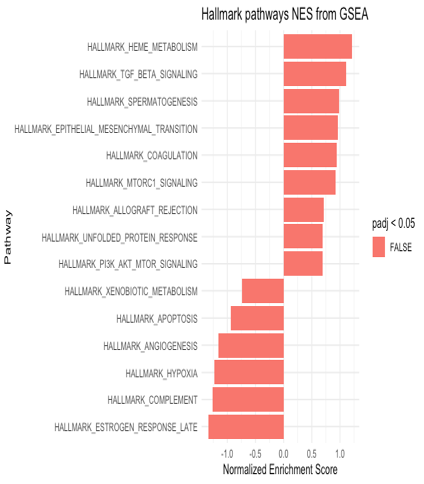

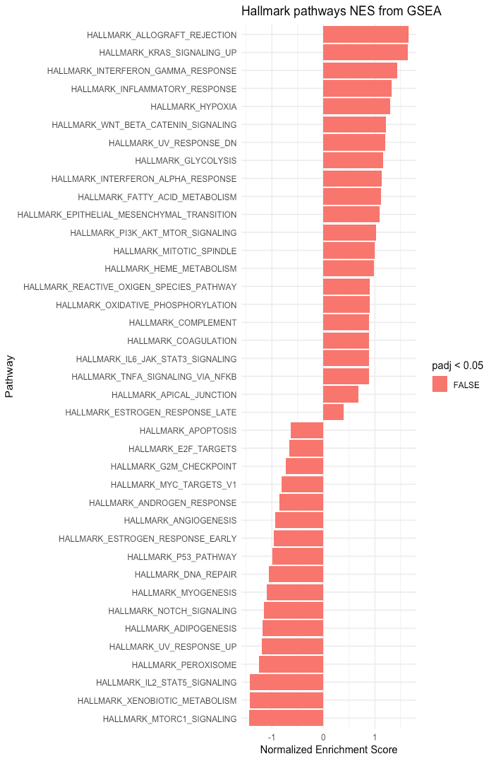

**A**

**B**

**C**

Supplementary Fig. 5. Hallmark identified common pathways from GSEA in persistently infected vs untreated. A) Hallmark identified common pathways from GSEA in TCCSUPPi vs EJ28Pi cells. B) Hallmark identified pathways from GSEA in TCCSUPPi cells. C) Hallmark identified pathways from GSEA in EJ28Pi cells. All the identified pathways are not significantly upregulated or downregulated (*p* <0.05) but are markedly dysregulated due to persistent NDV infection.

Supplementary Table 5 : Expression profiles of stress proteins and transcription factor genes

| **Gene symbol** | **Official gene name** | **log2FC** | **Adj *p* (FDR)** |
| --- | --- | --- | --- |
| **Stress proteins genes** |  |  |  |
| *HSPA1B* | Heat shock protein family A (Hsp70) member 1B | -7.4251934 | 0.01013638 |
| *HSPA4* | Heat shock protein family A (Hsp70) member 4 | -1.0595119 | 0.04753617 |
| *HSPB7* | Heat shock protein family B (small) member 7 | -1.4650371 | 0.01690584 |
| **Transcription factor genes** | |  |  |
| *HOXA13* | Homeobox A13 | -2.4218888 | 6.60E-06 |
| *HOXB13* | Homeobox B13 | -2.7894297 | 3.99E-06 |

Supplementary Table 6: Expression profiles of apoptosis, cell cycle and immune-related genes

| **Gene symbol** | **Official gene name** | **log2FC** | **Adj *p* (FDR)** |
| --- | --- | --- | --- |
| **Apoptosis** |  |  |  |
| *EI24* | Etoposide-induced gene 24 | -3.8818588 | 0.00132709 |
| *EMP2* | Epithelial membrane protein 2 | -3.4097884 | 0.04129703 |
| *TP53* | Tumour protein p53 | -3.2753604 | 9.01E-05 |
| *PTGES* | Prostaglandin E synthase | -5.4799260 | 0.00124882 |
| **Apoptosis & Immune response** | |  |  |
| *FASLG* | Fas ligand | -5.1558986 | 0.00446099 |
| *SP1* | Sp1 transcription factor | -5.1944248 | 3.47E-13 |
| *STAT6* | Signal transducer and activator of transcription 6 | -4.5315302 | 0.03987829 |
| **Cell cycle & growth** |  |  |  |
| *MNT* | MAX network transcriptional repressor | -4.0474029 | 0.00063625 |
| *NDRG4* | NDRG family member 4 | 2.701741 | 0.0473417 |
| *ZNF346* | Zinc finger protein 346 | 5.5467021 | 0.00196073 |
| *FGF14* | Fibroblast growth factor 14 | 5.3677443 | 0.00431774 |
| *FGF9* | Fibroblast growth factor 9 | 4.9654130 | 0.00852769 |

Supplementary Table 7 : Expression profiles of solute carrier and potassium voltage-gated channel genes

| **Gene symbol** | **Official gene name** | **log2FC** | **Adj *p* (FDR)** |
| --- | --- | --- | --- |
| **Solute carrier family** | |  |  |
| *SLC1A5* | Solute carrier family 1 member 5 | -2.4095596 | 1.95E-84 |
| *SLC43A3* | Solute carrier family 43 member 3 | -5.3068906 | 0.00418441 |
| *SLC48A1* | Solute carrier family 48 member 1 | -3.5350879 | 0.00024674 |
| **Potassium voltage-gated channel subfamily** | |  |  |
| *KCNJ10* | Potassium voltage-gated channel subfamily J member 10 | -6.4658741 | 1.60E-05 |
| *KCNJ12* | Potassium voltage-gated channel subfamily J member 12 | -5.639902 | 0.00074896 |
| *KCNJ5* | Potassium voltage-gated channel subfamily J member 5 | -6.3878306 | 2.19E-05 |

1.	Flicek P, Amode MR, Barrell D, Beal K, Billis K, Brent S, Carvalho-Silva D, Clapham P, Coates G, Fitzgerald S: **Ensembl 2014.** *Nucleic acids research* 2013, **42:**D749-D755.

2.	Pertea M, Kim D, Pertea GM, Leek JT, Salzberg SL: **Transcript-level expression analysis of RNA-seq experiments with HISAT, StringTie and Ballgown.** *Nature protocols* 2016, **11:**1650.

3.	Patro R, Duggal G, Love MI, Irizarry RA, Kingsford C: **Salmon provides fast and bias-aware quantification of transcript expression.** *Nature methods* 2017, **14:**417.

4.	Love MI, Huber W, Anders S: **Moderated estimation of fold change and dispersion for RNA-seq data with DESeq2.** *Genome biology* 2014, **15:**550.

5.	Team RC: **R: A language and environment for statistical computing.** 2013.
